# Patient-derived xenografts and single-cell sequencing identifies three subtypes of tumor-reactive lymphocytes in uveal melanoma metastases

**DOI:** 10.1101/2023.05.16.540908

**Authors:** Joakim Karlsson, Vasu R. Sah, Roger Olofsson Bagge, Irina Kuznetsova, Munir Iqbal, Samuel Alsén, Sofia Stenqvist, Alka Saxena, Lars Ny, Lisa M. Nilsson, Jonas A. Nilsson

## Abstract

Uveal melanoma (UM) is a rare melanoma originating in the eye’s uvea, with 50% of patients experiencing metastasis predominantly in the liver. In contrast to cutaneous melanoma, there is only a limited effectiveness of combined immune checkpoint therapies, and half of patients succumb to recurrent disease after two years. This study aimed to provide a path towards enhancing immunotherapy efficacy by identifying and functionally validating tumor-reactive T cells in liver metastases of patients with UM. We employed single-cell RNA sequencing of biopsies and tumor-infiltrating lymphocytes (TILs) to identify potential tumor-reactive T cells. Patient-derived xenograft (PDX) models of UM metastases were created from patients, and tumor sphere cultures were generated from these models for co-culture with autologous or MART1-specific HLA-matched allogenic TILs. Activated T cells were subjected to TCR sequencing, and the TCRs were matched to those found in single-cell sequencing data from biopsies, expanded TILs and in livers or spleens of PDX models injected with TILs. Our findings revealed that tumor-reactive T cells resided not only among activated and exhausted subsets of T cells, but also in a subset of cytotoxic effector cells. In conclusion, combining single-cell sequencing and functional analysis provides valuable insights into which T cells in UM may be useful for cell therapy amplification and marker selection.

## Background

Uveal melanoma (UM) is a rare form of melanoma and the most common primary malignancy of the eye^1^. It develops in the uvea of the eye, most often in the choroid and ciliary body, and more infrequently in the iris. Primary UM can be cured by brachytherapy or enucleation of the eye, but 50% of patients will develop metastasis^2^. The most common route for metastatic spread is the liver for reasons that are poorly understood. Patients with metastatic disease have a median survival of less than a year ^3^; however, ongoing clinical studies suggest that some treatments could prolong survival ^4^. Monotherapy immune checkpoint inhibitors (ICI) are markedly less effective in patients with UM^5,6^ than in those with metastatic cutaneous melanomas. However, combination treatments with PD-1/CTLA-4 inhibitors^7–9^ or PD-1/HDAC inhibitors^10,11^ have demonstrated longer overall survival than historic benchmark data ^3^. Tebentafusp, a T-cell engager suitable for patients with the HLA- A2 genotype^12,13^, is the first therapy to show a prolonged overall survival in a phase 3 randomized trial, increasing median survival from 16.0 months to 21.7 months ^14^. Another development is locoregional treatments, where a recent phase 3 trial demonstrated that isolated hepatic perfusion (IHP) triples hepatic progression-free survival^15^ compared with the best alternative care and historic benchmark data. Although overall survival data are not yet mature, retrospective data suggest a benefit with liver-directed therapy^16^. However, invariably, all patients progressed on both tebentafusp and IHP; therefore, more research is needed.

Adoptive cell therapy (ACT) with TILs or CAR-T cells has not been extensively studied in UM. Pilot data from a clinical trial ^17^ showed that TILs can cause responses in patients, and we have previously showed that HER2 CAR-T cells can eradicate UM in xenografts ^18^. There is, however, a lack of robust *ex vivo* screening models and very few patient-derived xenograft (PDX) mouse models from metastases ^19^ to use in studies to improve ACT. Part of the lack of success for ACT may be attributable to a lack of tumor-reactive lymphocytes, possibly owing to lower mutation burden in UM compared to cutaneous melanoma ^20^. However, both the clinical trial and previous analyses of TILs in UM liver metastases have suggested that at least some TILs are tumor reactive^17,21–23^.

Defining tumor-reactive T lymphocytes (TRLs) among TILs would enable the identification of candidate biomarkers for selective expansion or TCR transgenics in cell therapy experiments in mouse models and cell therapy trials. To decipher how immunotherapy can be made more effective, the aim of this study was to identify and functionally validate TRLs in metastases of patients with UM. To this end, we used paired single-cell RNA (scRNA) and TCR sequencing, as well as functional experiments using patient-derived xenograft (PDX) models and autologous TIL cultures.

## Results

We previously collected metastases of UM from patients in routine clinical care and two clinical trials. These were the SCANDIUM phase 3 randomized trial comparing IHP to best alternative care^24^ and the PEMDAC phase 2 single-arm trial where patients received a combination of the PD-1 inhibitor pembrolizumab and the HDAC inhibitor entinostat ^11^. Owing to the limited amount available of these biopsies, we prioritized one part of the biopsy for DNA/RNA preparation to sequence^22^, one part for transplantation into immunocompromised mice to generate PDX mouse models, one part for generation of TILs, one part for cryopreservation of finely minced tumor (flow cytometry and single-cell sequencing) and one part for formalin-fixed paraffin embedded blocks (FFPE) (**Fig. 1a**).

**Figure 1.**
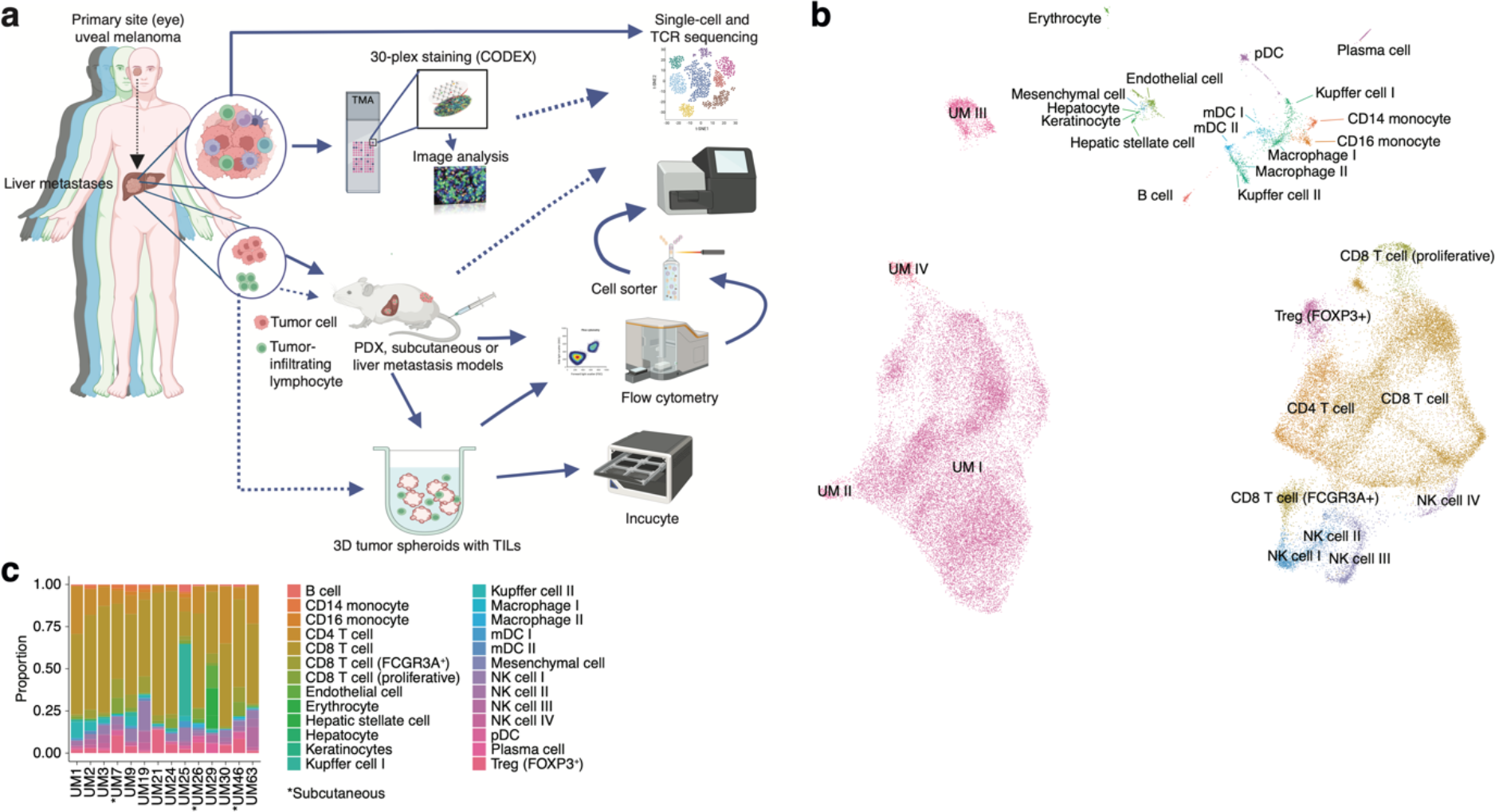
**a)** Schematic showing the study workflow. Tumors extracted from patient liver metastases were subjected to scRNA-seq, formalin fixation for immunohistochemical (IHC) analysis and PDX generation. **b)** UMAP dimensionality-reduced expression profiles of single cells in biopsies from 14 patients with UM. scRNA-seq data from all samples were integrated using FastMNN, Louvain clustering performed and clusters annotated using cell type marker genes compiled from the literature. Some cell types were annotated after additional subclustering (**Supplementary Fig. 1**a-f). **c)** Cluster proportions within each sample, for all cell types. Subcutaneous biopsies are indicated with asterisk, remaining samples being liver biopsies.

### Single-cell sequencing defines an atlas of T cell states in UM metastases

We obtained biopsies suitable for sequencing from 14 patients, two of which were subcutaneous and 12 were liver metastases. To identify and profile the T cells present in these samples, we performed paired scRNA and TCR sequencing, integrated the samples, and annotated the cell types. This has resulted in the largest single-cell atlas of metastatic UM to date, complementing prior efforts, including our own study of TILs from UM metastases ^21,22,25,26^.

Uniform Manifold Approximation and Projection (UMAP) dimensionality-reduced expression profiles revealed clusters containing melanoma, CD8^+^ or CD4^+^ T cells, NK cells, monocytes, and small clusters of B cells, endothelial cells, and other minor cell populations (**Fig. 1b-c**, **Supplementary Fig. 1**a-f). Clusters were generally well-mixed with respect to contributions from different samples, suggesting few patient-specific or batch effects after data integration (**Supplementary Fig. 1**g). The identity of cells labeled as UM was further confirmed by inferring copy number changes. This revealed characteristic aberrations seen in metastatic UM, including monosomy of chromosome 3 and gain of 8q, as well as subclonal variation within patients (**Supplementary Fig. 1**h).

Further sub-clustering of CD8^+^ T cells showed the presence of 12 different groups (**Fig. 2a-b** and **Supplementary Fig. 1**i-k), all characterized by the differential expression of immunological marker genes (**Fig. 2c-d** and **Supplementary Table 1**). The clusters can be divided into two axes. One of these is characterized by expression of genes associated with late activation and exhaustion, such as *PDCD1*, *HAVCR2*, *TNFRSF9* and *HLA-DRA*. Additional expression of *ITGAE* (CD103) suggests that these clusters contain tissue-resident memory cells^27^. Multiplex immunofluorescence showed that PD-1- and ICOS-expressing T cells were in close proximity to the tumor cells (**Fig. 2e**). The other axis expressed genes associated with naïve/memory-like or early activated phenotypes, such as *IL7R*, *TCF7*, *CCR7* and *NR4A1* (**Fig. 2d**).

**Figure 2.**
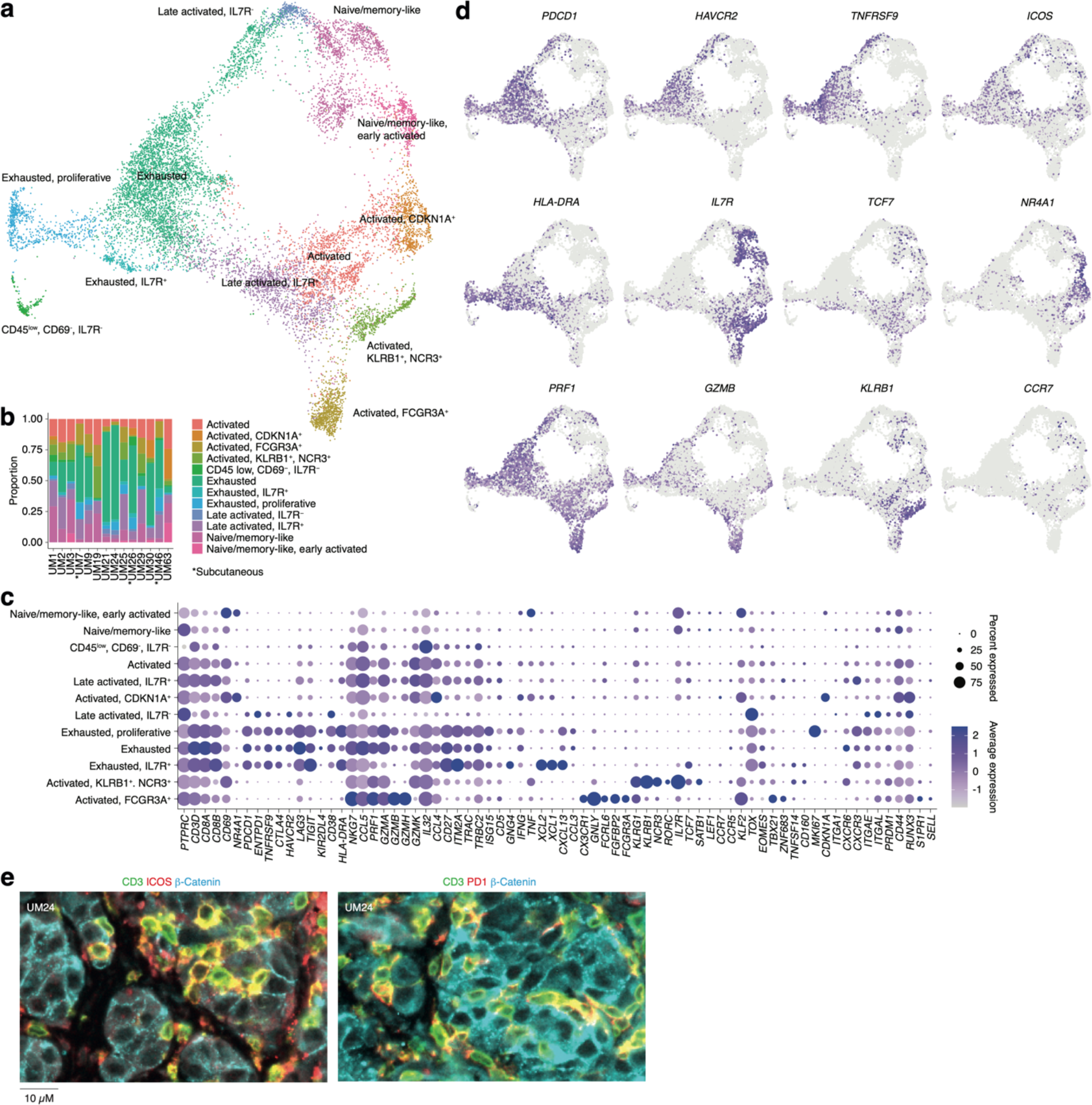
**a)** UMAP dimensionality reduction and more detailed re-annotation of the CD8^+^ T cell subsets from Fig. 1b, using the marker genes shown in (c). **b)** CD8^+^ T cell cluster proportions within each sample. **c)** Expression levels of markers used for CD8^+^ T cell subset annotation. **d)** Expression of marker genes signifying early (*NR4A1*) activation, naïve/memory-like phenotypes (*TCF7*, *IL7R*), late activation (*HLA-DRA*, *PDCD1, ICOS*) and progression towards exhaustion / dysfunction (*LAG3*, *TIGIT*, *HAVCR2*). Additional statistically identified genes differing between clusters can be found in **Supplementary Table 1**. **e)** Multiplex IHC staining for T cell (CD3, ICOS, PD-1) and cancer marker genes (β-catenin) in a patient biopsy (UM24).

Individual clusters within these axes mostly correspond to intermediate phenotypes that progress from activation to exhaustion. However, a few seem to represent distinct states. Two of these were within the *IL7*^+^ axis. Of these, one expressed *KLRB1* and *NCR3* and the other expressed NK-associated markers, such as *FCGR3A*, *FCRL6* and *NKG7* (**Fig. 2c-d**). Although both clusters expressed genes related to cytotoxicity, only the latter expressed *GZMB* and *GZMH*. Other markers associated with this group are *CX3CR1* and *FGFBP2*. Previous studies labeled these cells as cytotoxic NK-like T cells^28,29^.

In summary, this map reveals the cell types present in liver and subcutaneous metastases of UM and divides infiltrating T cells into two major phenotypic groups, characterized by the mutually exclusive expression of *HAVCR2* and *IL7R*, but also reveals distinct cytotoxic subsets expressing NK-associated markers.

### TILs from UM metastases can kill their cognate tumors in PDX models

Given the presence of activated T cells in tumors, we investigated whether these could eradicate metastatic UM in ACT experiments using PDX models. These models are useful personalized models for studying tumor biology and more recently, tumor immunology^30^. However, the biopsies we received were generally very small, and the initial take rate using the same single-cell dispersion method used for cutaneous melanoma metastases^30^ was unsatisfactory. Therefore, we transplanted small tumor pieces or used splenic injections, which led to successful PDX generation for some patients. Ten of these are presented in this study. All the PDX tumors exhibited confirmatory markers of melanoma, such as SOX10, MELAN-A (MART1), HMB45 or S100B. Sanger sequencing further revealed mutually exclusive mutations in the UM oncogenes *GNAQ*, *GNA11*, *CYSLTR2* and *PLCB4*, as found in biopsies^22^. (**Fig. 3a** and **Supplementary Fig. 2**). The UM tumors grew at a slower rate compared to our experience with cutaneous melanoma, which has a take rate of 95% and where a model is usually established within 3-12 weeks (**Fig. 3b**)^31^.

**Figure 3.**
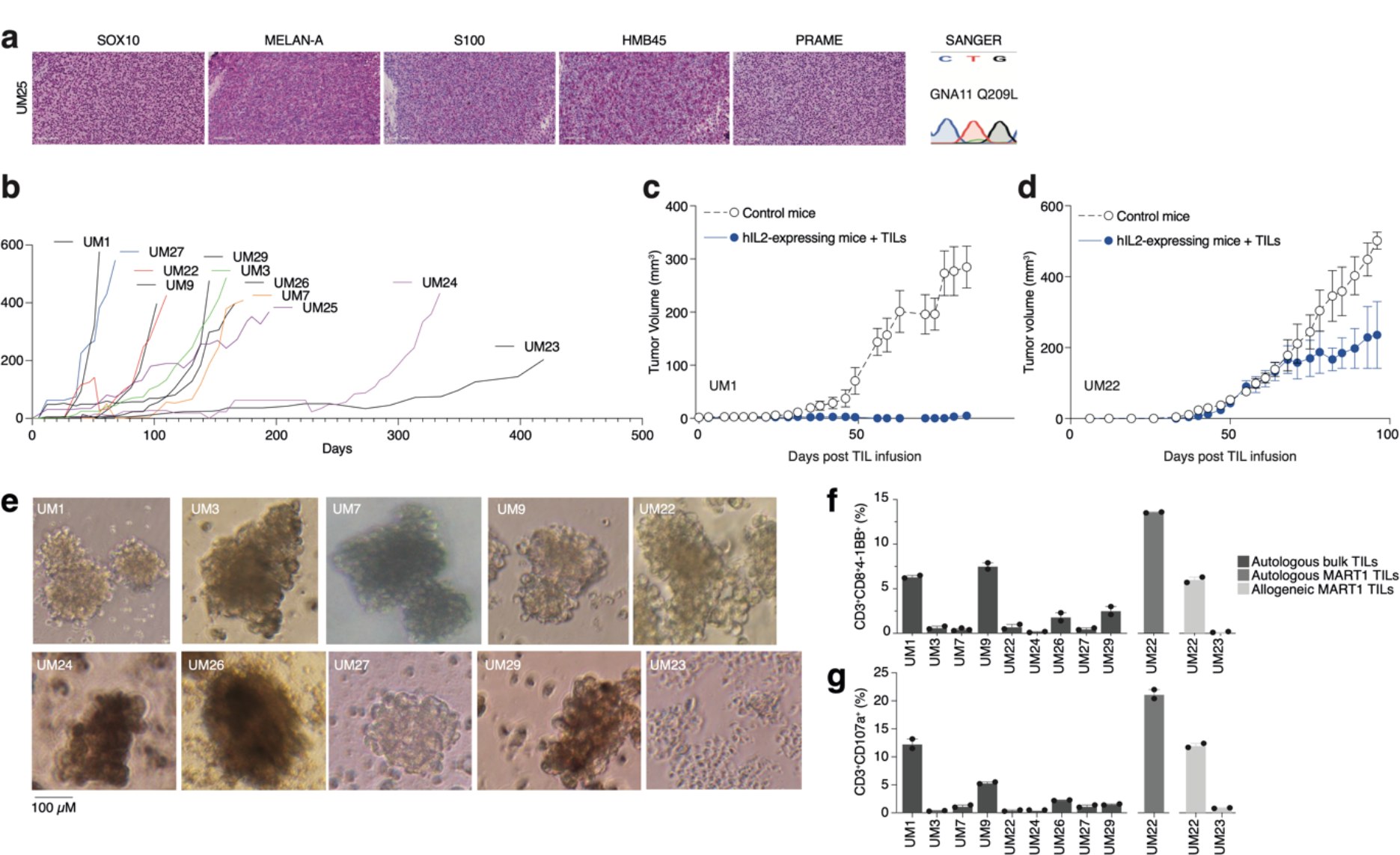
**a)** IHC staining of HMB45, SOX10, S100, MELAN-A (MART1) and PRAME expression in one PDX (See also **Supplementary Fig. 2** for additional PDX models), as well as Sanger sequencing to verify mutation of *GNA11* at Q209L. Color reactions were with magenta. **b)** Tumor growth curves of established PDX models. **c-d**) TIL therapy experiments in PDX models. UM1 and UM22 were transplanted subcutaneously into NOG or hIL2-NOG mice. When tumors were palpable, 20 million autologous TILs were injected in the tail vein. Tumor growth was monitored by a caliper. Tumor sizes are average size ± SEM. **e)** Images of tumor spheroids created (scale bar represents 100uM). UM23 was unable to form spheres. **f-g)** Activation (41BB^+^) and degranulation (CD107a^+^) status for spheres and TIL co-cultures, (f) and (g) respectively, using *n* = 2 or 3 replicates.

We have previously developed a method to study ACT in PDX models, termed PDXv2^18,32^. Survival, expansion, and activity of TILs and CAR-T cells were improved in this model using human IL2-expressing transgenic NOD SCID IL2 receptor gamma knockout mice (hIL2-NOG)^18,32–35^. Tumor eradication by TILs correlates with immunotherapy response in patients with melanoma^32,36^. We conducted a PDXv2 experiment using PDX models from two patients with UM. The injection of autologous TILs into hIL2-NOG mice carrying UM1 resulted in tumor rejection, whereas UM22 tumors could not be eradicated (**Fig. 3c-d**). This constitutes proof of concept that TILs from UM metastases can kill autologous tumors in a PDXv2 model.

### TILs contain subsets capable of e*x vivo* expansion and killing in 3D tumor sphere cultures

To gain further insight into the capacity of T cells in UM biopsies to kill tumors, we next investigated the tumor reactivity of *ex vivo* expanded TILs on autologous UM cells. Since most UM PDX models did not grow fast enough to routinely conduct ACT experiments, we used the tumors grown in the mice to generate 3D tumor sphere cultures ^37,38^ of high viability for TIL experiments^39,40^. We generated tumor sphere cultures from ten PDX models and expanded their autologous TILs (**Fig. 3e**). When tumor spheres and TILs were co-cultured, TILs from patients UM1 and UM9 were activated the most (surface 4-1BB^+^) and also degranulated (surface CD107a^+^; **Fig. 3f-g**). These TILs also bound to the tumor spheres, as assessed by labeling tumor spheres and TILs with different vital dyes and then imaging with a real-time cell culture imager (**Supplementary Fig. 3**a-b). The UM22 TILs were hardly able to get activated by their autologous tumor spheres (**Fig. 3b**), which likely explains their lack of activity in the PDX ACT experiment (**Fig. 3d**). However, since UM22 is capable of presenting MART1 on HLA-A2, and since we previously identified a minor population of MART1-reactive TILs in this biopsy^41^, we decided to expand and enrich this subset using MART1 peptide HLA-A2 dextramers. When using these cells, we were able to achieve potent activation and degranulation (**Fig. 3f-g**). This confirms that T cells extracted from biopsies can contain subsets capable of recognizing and killing tumors, which can be expanded *ex vivo*.

### Determining the phenotypes of TRLs in biopsies using scRNA and TCR sequencing

To gain clues on how to identify reactive T cells from biopsies through their transcriptomic phenotypes, we co-cultured UM1 and UM9 TILs with their autologous tumors and sorted 4-1BB activated T cells by cell sorting. We prepared RNA from these cells, and as a control, from MART1-specific allogenic UM46 TILs and sequenced their TCRs. Then, using the single-cell atlas built from biopsies, we mapped those TCRs back to T cell subsets from their original tumors. This showed that those TRLs predominantly emanated from exhausted and late-activated cells (**Fig. 4a-b**). UM9 also contained a sizeable amount of TRLs in the NK-like/cytolytic cluster (**Fig. 4b**). The lack of potential TRL clones identified in the *TCF7*^+^ axis of the CD8^+^ T cell atlas from biopsies is also compatible with a previous report showing that *TCF7*^+^ T cells in melanoma tend to be nonreactive bystanders^28^.

**Figure 4.**
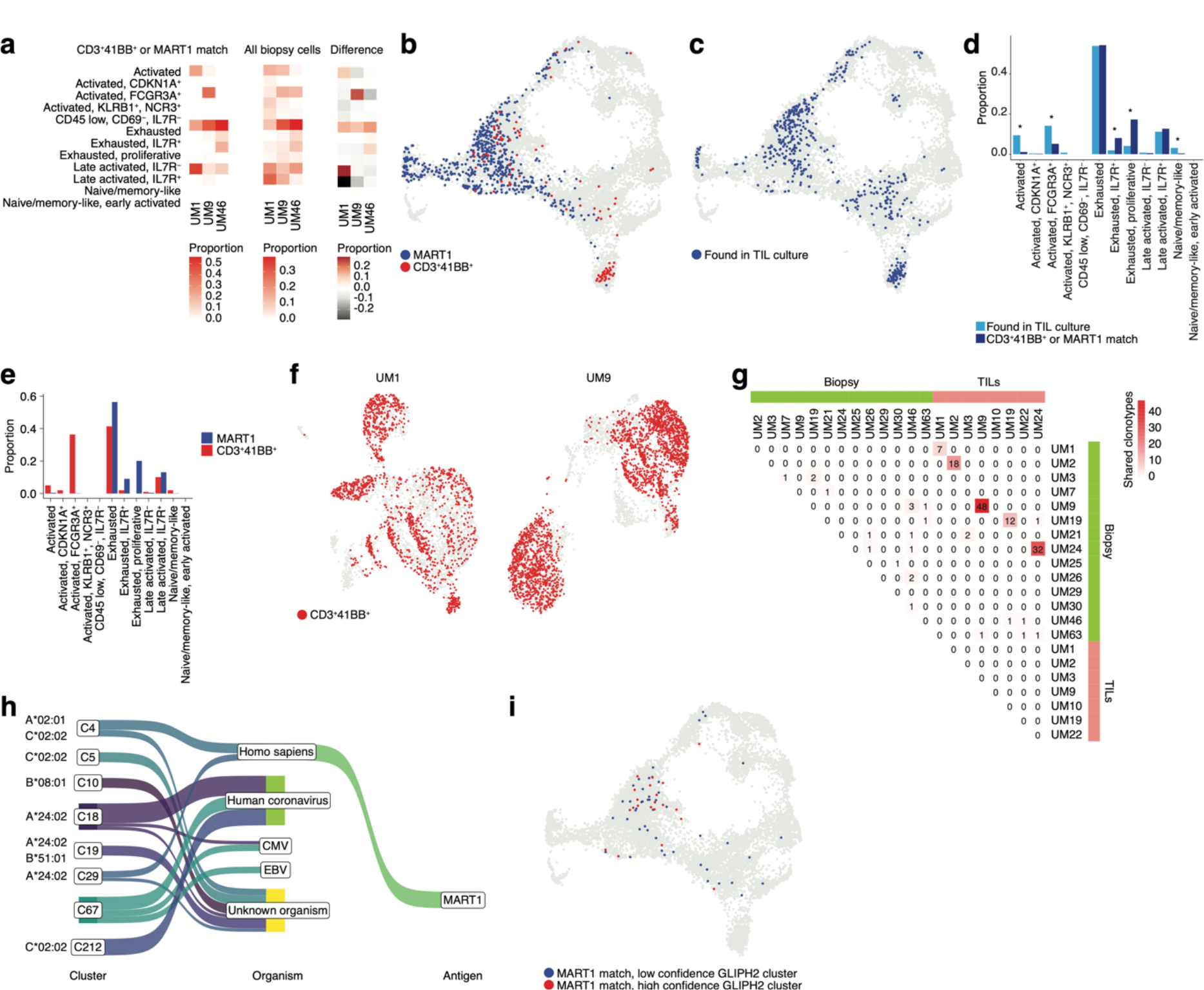
**a)** Proportions of cells from each sample that match clonotypes found in experimentally identified 4- 1BB^+^ and MART1-reactive T cells and which are present in a given biopsy CD8^+^ T cell cluster. The difference between matching and non-matching cells in each cluster is shown on the right, highlighting subsets that are enriched among the reactive T cells. **b)** Biopsy CD8^+^ T cells with clonotypes matching identified reactive T cells, highlighted in the UMAP representation. **c)** As in (b), but highlighting cells with clonotypes shared between biopsies and TILs. **d)** Proportions of biopsy CD8^+^ T cells in each cluster matching either clonotypes from TIL cultures or experimentally identified reactive T cells. Statistical differences in frequency were determined using binomial tests between the frequency of the latter in each cluster relative to the background frequency of all TIL clonotypes present in the same cluster. Frequencies were calculated as number of cells from a given category in a given cluster divided by the total number of cells from that category, where category refers to either TILs or 4- 1BB^+^/MART1-reactive cells. *p*-values were adjusted for multiple testing using Bonferroni correction. **e)** Distributions of cells matching the two different categories of experimentally identified reactive cells among biopsy CD8^+^ T cells clusters. **f)** Cells from scRNA-seq of TIL cultures that have clonotypes matching experimentally identified reactive cells. **g)** Shared clonotypes among all biopsy and TIL samples. **h)** GLIPH2 clusters of significantly similar and HLA-restricted clonotypes^43^, mapped to known antigens in public databases using TCRMatch^42^, based on CDR3β sequences. **i)** Biopsy CD8^+^ T cells with clonotypes in either high or low confidence GLIPH2 clusters that match MART1 motifs in public databases.

To evaluate which T cells in biopsies were reflective of those capable of generating TIL cultures, we further mapped our previously published TCR sequencing data of UM TILs to the biopsy data (**Fig. 4c**) to compare them to the clones matching experimentally identified TRLs in the biopsy (**Fig. 4b-d**, **Supplementary Fig. 4**a-b and **Supplementary Table 2**). Cells from the TIL dataset tended to correspond to biopsy T cells with exhausted and late-activated profiles, similar to the experimentally identified TRLs. This might be expected since both have undergone expansion, which might favor T cells in certain phenotypic states. However, TRLs were significantly over- and underrepresented in some phenotypic clusters compared to TILs in general. Overrepresented clusters included proliferative exhausted and *IL7R*^+^ exhausted T-cells. The underrepresented ones were activated (non-exhausted), naïve/memory-like, and *FCGR3A*^+^ NK-like T cells. The distribution of TRLs across the clusters was also different for cells sorted by 4-1BB-positivity versus MART1-recognition. MART1-reactive cells were more likely to be found in exhausted and late-activated clusters than 4-1BB^+^ cells, which tended to map to earlier activation states as well to the NK-like cluster (**Fig. 4e**). The latter may be partly due to their prevalence in UM9 (**Supplementary Fig. 4**c).

Mapping TCRs from enriched 4-1BB^+^ cells to TILs in corresponding patients from previously published TCR sequencing data revealed that a large fraction of TILs in these cultures was among the identified TRL clones (**Fig. 4f**). In 6 of the 8 previously published TIL cultures, we were able to recover and sequence the corresponding cryopreserved biopsy of that patient. Five of the six cultures had many TCRs that overlapped between TILs and biopsies (**Fig. 4g** and **Supplementary Fig. 4**a-b). UM3 did not have any matching TCRs between the biopsies and expanded TIL cultures. No overlap was observed between samples from different patients with the 4- 1BB^+^/MART1^+^-selected TCRs (**Supplementary Fig. 4**d). This is reasonable given the different HLA genotypes of these patients. In general, very few clonotypes were shared between samples from different patients (**Supplementary Fig. 4**e). In summary, experimentally identified TRLs mapped back to biopsies were overrepresented in proliferative exhausted and *IL7R*^+^ exhausted T cells, and these T cells could also be identified in sequenced cultures of expanded TILs.

### Identifying potential antigens recognized by TRLs

To understand potential antigens recognized by a given TCR, we searched for matches in public databases of known TCRβ chains, based on CDR3β motifs, using TCRMatch^42^. The results confirmed that a number of TCRs matched known MART1-reactive clonotypes but also revealed matching TCRs that might recognize other melanoma antigens, such as PMEL and MAGEA1 (**Supplementary Fig. 4**f-g, **h**). Overall, a large number of TCRs also matched clonotypes recognizing viral antigens such as influenza, EBV, and human coronaviruses. However, an even greater number lacked known matches, suggesting they might recognize tumor-specific antigens (neoantigens).

However, public databases are currently limited. As an alternative approach to nominate tumor-reactive clonotypes, we used an algorithm designed to predict clusters of TCRs that are sufficiently similar and HLA-restricted to potentially recognize the same antigens. GLIPH2^43^ nominated a few high-confidence clusters of similar TCR sequences based on the biopsy scRNA-seq data (**Fig. 4h** and **Supplementary Table 3**). To further test the hypothesis that each cluster recognizes the same antigen, we mapped the CDR3β motifs of these sequences to the same public databases. Interestingly, the highest-ranked cluster was overrepresented for sequences matching MART1, and this cluster also exhibited significant restriction towards HLA-A*02:01, on which MART1 is known to be presented. The remaining high-confidence clusters mainly matched antigens of viral origin or lacked matches to known antigens (**Fig. 4h**).

Similar to the experimentally determined MART1-reactive T cells, the bioinformatically determined cluster of potential MART1-recognizing clones mostly belonged to the exhausted CD8^+^ T cell cluster (**Fig. 4i, Supplementary Fig. 4i**). This supports the finding that TRLs found in biopsies may have predominantly exhausted profiles. In addition, a high-confidence group of clonotypes co-localized to the NK-like cluster was nominated by GLIPH2, although no sequences from this cluster matched the public TCR-antigen pairs (**Fig. 4h**, **Supplementary Fig. 4**i). Given that the cells from this cluster were experimentally identified as 4-1BB^+^ upon tumor co-culture (**Fig. 4b**), one might speculate that they could recognize unknown tumor antigens. Taken together, these findings suggest that both bioinformatically inferred TRLs from biopsies and *ex vivo* expanded TRLs tend to have exhausted phenotypes. Among these, T cells recognize known melanoma antigens such as MART1, PMEL, and MAGEA1, as well as unknown sequences that might be neoantigens.

### TRLs home to the tumor of PDX mice bearing liver metastases of UM

TIL cultures contain both TRLs, virus-specific T cells, and bystander T cells that grow during IL2-containing culture conditions. We had previously found that TILs can home to subcutaneously growing skin melanoma metastases ^18,32^ but we were interested in determining whether they could identify and home to UM growing in the liver. The PDX models used to generate sphere cultures were prepared by subcutaneous transplantation. To instead generate a liver metastasis model, we performed splenic injection of UM22 into NOG mice^44^. Tumor cells forming liver metastases were further transplanted via tail vein injection into hIL2-NOG mice. After verifying tumors by ultrasound, autologous bulk or MART1-selected UM22 TILs were intravenously injected for TIL therapy studies (**Fig. 5a**). The mice were harvested three weeks later, and tumors recovered from the liver were analyzed by flow cytometry, IHC, single-cell RNA and TCR sequencing. As expected from **Fig. 3f-g**, MART1-selected TILs had a higher degree of activation than unselected bulk UM22 TILs (**Fig. 5b**), despite similar numbers of TILs present in the livers. There were no visible differences in the size of the liver tumors between mice receiving unselected or MART1-selected UM22 TILs. This may be due to T-cell exclusion, since most TILs colonized the rim of the tumor (**Fig. 5c**). More MART1-specific TILs than unselected TILs were able to get activated, as assessed by the activation markers 4-1BB and CD69 (**Fig. 5b** and **d-e**). This shows that T cells recognizing a tumor antigen can successfully migrate to liver metastases of UM in mice.

**Figure 5.**
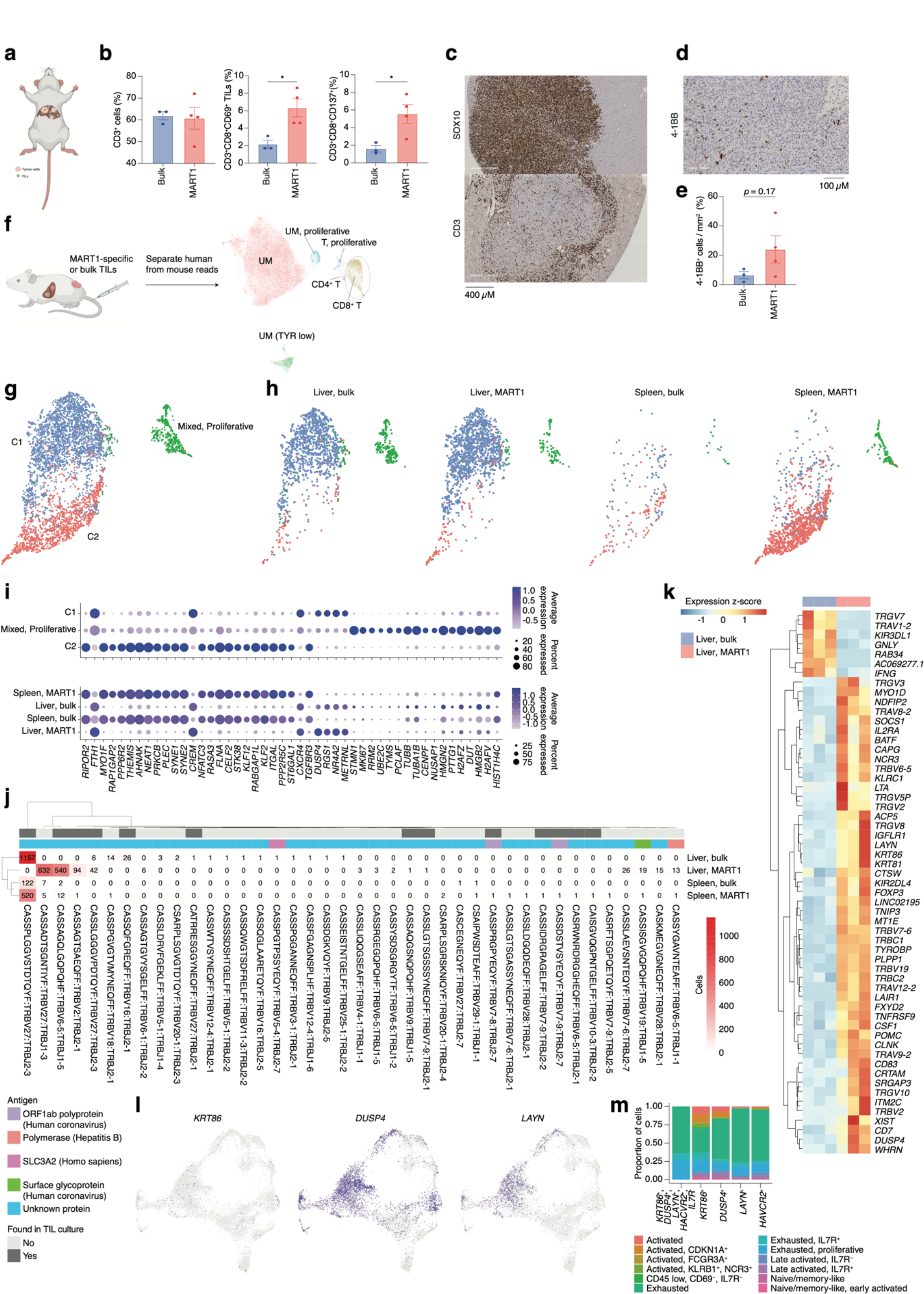
**a)** Bulk unsorted TILs or MART1 selected TILs were injected into hIL2-NOG mice carrying liver tumors from patient UM22. **b)** Flow Cytometry analysis of single cell suspension liver metastasis, comparing treatment of UM22 TILs and UM22 MART1-specific TILs for CD3^+^, CD3^+^CD8^+^CD69^+^ or CD3^+^CD8^+^CD137^+^. **c)** IHC with diaminobenzidine (DAB) showing tumor (SOX10) and TILs (CD3) within a liver metastasis, **d-e)** corresponding analysis of 4-1BB^+^ TILs in a section (d) and in image analysis comparing both treatments (e). Statistical tests in b and e were unpaired two-tailed *t*-tests, assuming equal variance. *: *p* < 0.05; **: *p* < 0.01. **f)** Samples of tumor and TILs from the liver and spleen, respectively, were sequenced with scRNA-seq. *n* = 3 biological replicates were performed for each group of liver samples, and *n* = 2 for spleen samples (out of which one spleen sample for MART1-selected TILs was the pooled material of two independent mice). Sequencing reads mapping to human and mouse were separated with XenoCell^66^, after which cells were clustered and annotated as described in **Methods**. **g)** Subclustering of CD8^+^ T cells identified three overall clusters, one of represented a mixed profile of the other two, but with marked cell cycle activity. **h)** Contributions of the different experimental conditions to each CD8^+^ T cell cluster. **i)** Markers distinguishing CD8^+^ T cell clusters, identified using the FindAllMarkers function of Seurat (the union of the top 25 genes per condition, ranked by log2 fold change). Expression per experimental condition is shown below. **j)** All TCRβ chains identified in each experimental condition. Subsets found in TIL culture scRNA-seq data are highlighted, as are any matches to antigens in public databases. **k)** Differentially expressed genes between bulk TIL mixtures or MART1-selected TILs present in the livers of mice. A pseudo-bulk approach was used, summing read counts across all cells within a given replicate, and statistical testing performed with DESeq2^81^. Genes with *q* < 0.05 after Benjamini-Hochberg correction were considered significant. **l-m)** Expression of *KRT86*, *DUSP4* and *LAYN* in biopsy CD8^+^ T cells (l) and the phenotypic clusters they are members of (m).

### Determining the phenotypes of TRLs in PDX liver metastases using scRNA and TCR sequencing

Next, we profiled the livers and spleens of mice injected with unselected bulk or MART1-selected UM22 TILs by scRNA and TCR sequencing (**Fig. 5f**, **Supplementary Fig. 5**a). After bioinformatically separating human- from mouse-derived cells and integrating the data, clearly distinguishable clusters emerged containing UM cells, all characterized by the expression of the differentiation genes *TYR*, *MLANA* and *PMEL*. Their designation as UM cells was further confirmed by inference of their copy number profiles (**Supplementary Fig. 5**b). As expected, all UM cells resided in the liver (**Supplementary Fig. 5**a), despite the mice being injected with UM cells via the tail vein. The TILs discovered in these samples constituted three clusters: C1, C2, and a proliferative cluster with a mixed set of cells from the two main clusters (**Fig. 5g**). Compared with CD8^+^ T cell clusters from the biopsies, the clusters found in mice did not separate with respect to the same set of markers. Rather, all clusters expressed markers compatible with late-activated or exhausted cytotoxic cells to some extent (**Supplementary Fig. 5**c). These TILs were detectable in both the liver and the spleen, although there was a notable correspondence between, on one hand, liver TILs and C1, and on the other, spleen TILs and C2 (**Fig. 5g-i**).

Investigating the TCR clonotypes represented in the four experimental conditions revealed several clones unique to or overrepresented in the liver biopsies of mice with MART1-selected T cells, as compared to unselected TILs or spleen samples (**Fig. 5j**). Given that these clones have been sorted to enrich for reactivity towards an antigen known to be presented by UM22 cells, and that they have successfully migrated to the liver tumor, this subset is likely to be strongly enriched for TRLs. Among these, some were also found in previous scRNA-seq data from expanded UM22 TILs (**Supplementary Fig. 5**d-i).

Differential expression analysis between this TRL subset and unselected TILs found in the livers of PDX mice revealed statistically significant genes that may be potential markers of the tumor-reactive phenotype (**Fig. 5k, Supplementary** Fig. 5j**, Supplementary Table 4**). This included the activation marker *TNFRFS9* (4-1BB) as well as *KIR2DL4*, which has been found to be expressed in certain exhausted subsets of CD8^+^ T cells (**Fig. 2c**)^28^. Interestingly, this subset also bears a striking resemblance to a recently described group of TIM-3^+^IL7R^-^ tumor-reactive tissue-resident memory cells in lung cancer, further characterized by the specific expression of *KRT86*, *DUSP4* and *LAYN* (**Supplementary Fig. 6**)^45^. This set was enriched in patients responding to anti-PD1 therapy^45^. Cells expressing this marker combination were also present in biopsy samples, predominantly in exhausted clusters (**Fig. 5l-m**), as well as in TIL cultures from UM22 (**Supplementary Fig. 6**b). This might suggest avenues for enriching TRLs from biopsies for use with *ex vivo* expansion and ACT based on their gene expression profiles.

### CD8 cells diminish after tumor eradication in PDX models

Although the UM22 PDX model demonstrated signs of TIL homing and activation, TILs were unable to cure the mice from the tumor, neither when grown subcutaneously (**Fig. 3d**) nor in the liver (**Fig. 5c**). We therefore developed a liver metastasis model of UM1, which responds to TILs in a subcutaneous PDX model (**Fig. 3d**). Injection of TILs into hIL2-NOG mice bearing UM1 liver metastases resulted in a complete clearance or tumors as assessed macroscopically and by ultrasound imaging. To characterize the cellular composition of T cells in the livers and spleens of the PDX mice treated with TILs, we performed single-cell RNA and TCR sequencing. Clustering analysis identified 15 clusters of human cells that were annotated by known marker genes defined by differential expression analysis and supported by literature (**Fig. 6b-c, Supplementary** Fig. 6c-e**, Supplementary Table 5 and 6**). Notably, and at variance with the UM22 model, none of the clusters contained any melanoma cells, confirming that the TILs had eradicated the tumor.

**Figure 6.**
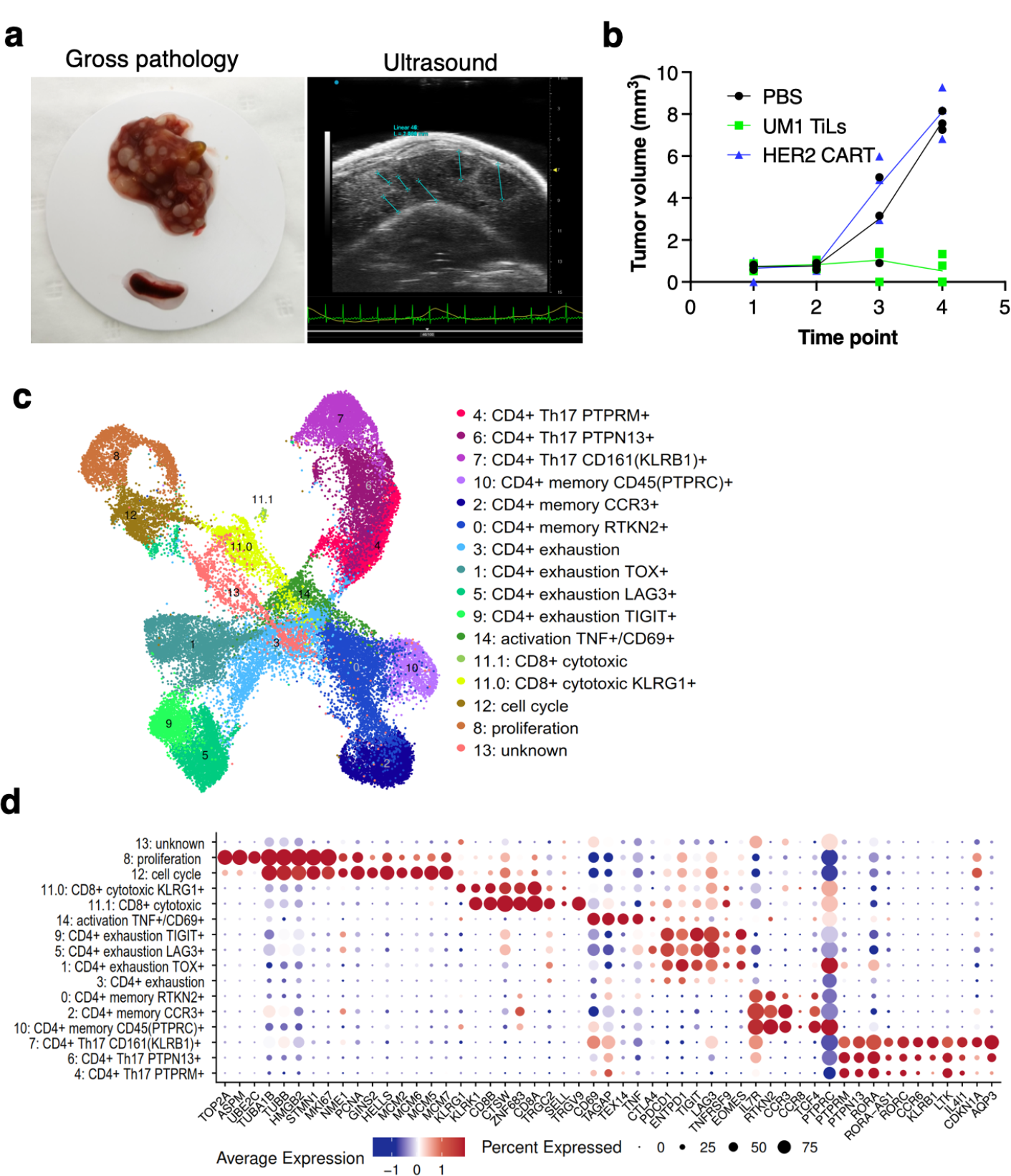
**a)** Establishment of another uveal melanoma liver metastasis model. UM1 PDX cells were injected in the tail vein after having been serially transplanted in spleen followed by harvesting from liver. Ultrasound confirmed growth in liver before injection of autologous UM1 TILs or HER2 CAR-T cells as controls. **b**) Response to TILs as assessed by ultrasound monitoring. **c**) UMAP of T cells in the UM1 liver metastatic model, showing 15 different cell populations. A list of marker genes was used to annotate clusters. The marker genes list was compiled from differential expression analysis and literature. **d**) Dot plot showing an average expression of marker genes and detection rate of cells in which the marker gene is detected across 15 cell populations.

We compared number of cells coming from liver and spleen samples. A total of 20,418 cells were found in liver and 18,208 from spleen (**Supplementary Fig. 7**a-b). Strikingly, even though the TILs injected were primarily CD8 positive, the T cells remaining in liver and spleen after tumor eradication were primarily CD4+ positive cells (**Supplementary Fig. 7**b-c). Most CD8 cells resided in Cluster 11 with fewer cells also in clusters 8 and 12 due to expression of cell cycle genes. Re-clustering of cluster 11 resulted in two subpopulations (11.0 and 11.1) annotated as CD8+ cytotoxic expressing killer cell lectin like receptor G1 (*KLRG1*) and CD8+ cytotoxic expressing higher levels of *CD8A* and *CD8B*. (**Supplementary Fig. 7**d). These cells also expressed higher levels of *CTLA4, TIGIT* and *TNFRSF9* (encoding activation marker CD137/4-1BB) and the ψ8Tcell markers *TRGC2/TRGV9*. Cluster 8 expressed high levels of Ki-67 (*MKI67*) assembly factor for spindle microtubules (ASPM), stathmin 1 (*STMN1*), DNA topoisomerase II alpha (*TOP2A*), TPX2 microtubule nucleation factor (*TPX2*) corresponding to the proliferative cellular state. Cluster 12 expressed minichromosome maintenance complex component 2, 6, 7 (*MCM2, MCM6, MCM5, MCM7*), GINS complex subunit 2 (*GINS2*), helicase, lymphoid specific (*HELLS*) that are associated with the proliferation and cell cycle state.

Next, we investigated the CD4+ cells and found that clusters 1, 3, 5, 9 expressed programmed cell death 1 (*PD-1*, also, known as *PDCD1*), lymphocyte activation gene 3 (*LAG3*), cytotoxic T lymphocyte-associated protein 4 (*CTLA-4*), T cell immunoreceptor with immunoglobulin and ITIM domains (*TIGIT*), thymocyte selection associated high mobility group box (*TOX*) and eomesodermin (*EOMES*) marker genes that are associated with a dysfunctional or exhausted state and was annotated as a CD4+ exhaustion TOX+, CD4+ exhaustion PTCD1+, CD4+ exhaustion LAG3+ and CD4+ exhaustion TIGIT+ respectively. Three adjacent clusters 0, 2, 10 expressed interleukin 7 receptor (*IL7R*), rhotekin 2 (*RTKN2*), C-C motif chemokine receptor 3 (*CCR3*) and protein tyrosine phosphatase receptor type C (PTPRC, also known as CD45) marker genes corresponding to a memory T cells phenotype and were annotated as a CD4+ memory RTKN2+, CD4+ memory CCR3+ and CD4+ memory PTPRC+ respectively. More liver cells contributed to the CD4+ exhaustion cluster whereas the CD4+ memory cluster was primarily made up of spleen cells (**Supplementary Fig. 7**b). Finally, three adjacent clusters 7, 6, 4 were associated with a CD4+ Th17 phenotype that play important role in the inflammatory response expressing the transcription factor retinoic acid orphan receptor (ROR)γt (*RORC*), chemokine CC receptor 6 (*CCR6*), a killer cell lectin like receptor B1 (*KLRB1*, also known as *CD161*), interleukin 4 induced 1 (*IL4I1*), cyclin dependent kinase inhibitor 1A (*CDKN1A*), aquaporin 3 (*AQP3*) genes (**Supplementary Fig. 7**e) and annotated as a CD4+ Th17 CD161+ (KLRB1), CD4+ Th17 PTPN13+ and CD4+ Th17 PTPRM+ respectively.

Finally, we investigated the TCR clonotype abundance and visualized top ten the most abundant clonotypes (**Fig. 7a**). Clonotypes 302, 11155 and 21797 were the most abundant, and as expected, they were present in CD4 + cells. Clonotypes 302 mapped to the CD4+ exhaustion site, 111555 to the CD4+ memory and 21797 to the CD4+ Th17 cluster (**Fig. 7b**). To investigate if there were any differences in gene expression between these clonally expanded T cells in liver versus spleen, we performed pseudobulk analysis. Genes such as *FASLG*, *ICOS*, *TIGIT*, *IL2RA* that were higher in T cells residing in liver are known to be expressed during activation and exhaustion. (**Fig. 7c**). We also mapped where TRLs are residing by extracting the top ten T cell Receptor Beta (TRB) sequences obtained from the CD137 sorted UM1 TILs (**Fig.3f-g** and **Fig.4).** We identified a few of these in the PDX data and mapped them on the UMAP (**Fig. 7d**). TRLs were residing on the CD4+ Th17 and proliferation/cell cycle clusters.

**Figure 7.**
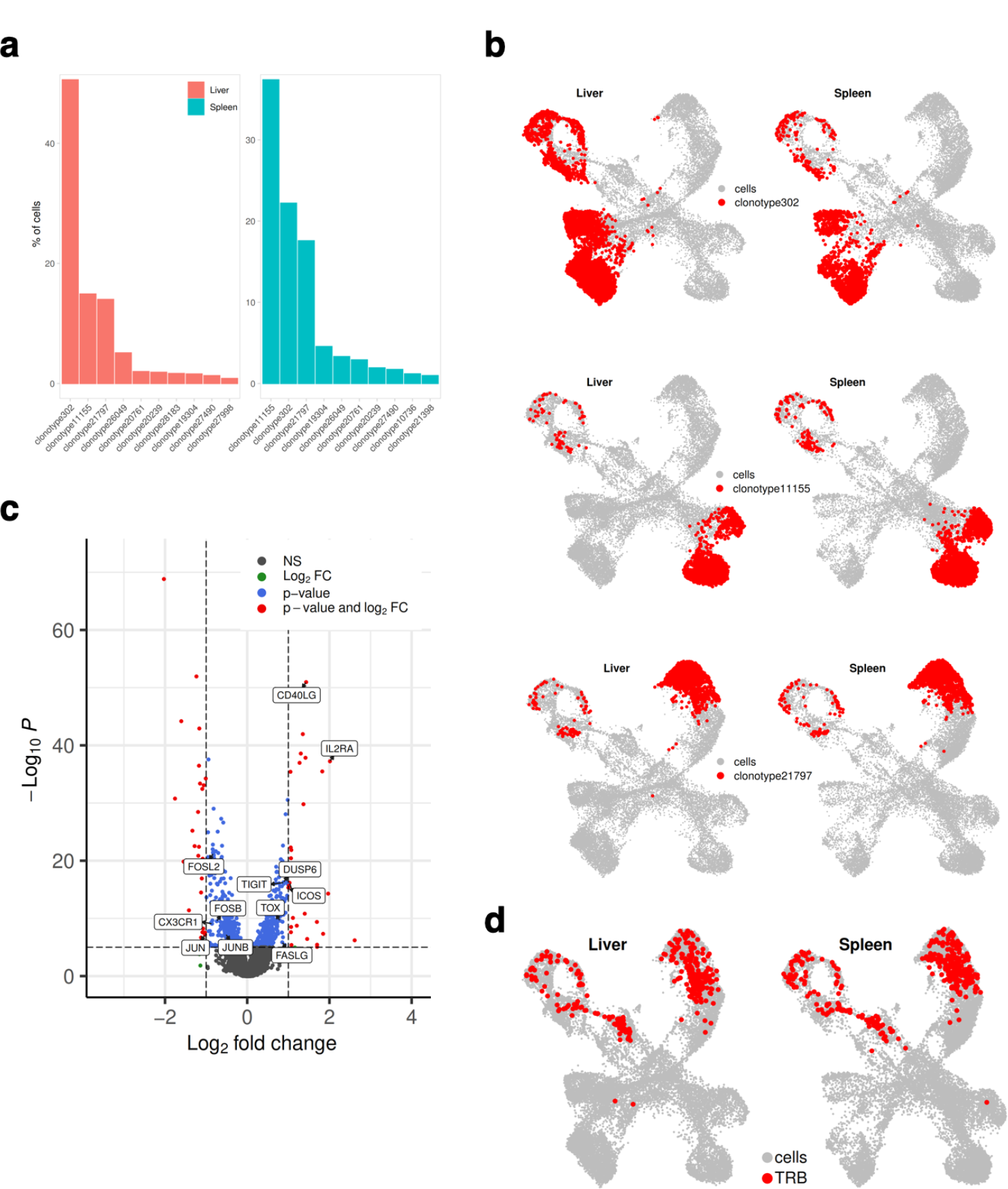
**a)** Bar plot showing top 10 the most abundant clonotypes for liver and spleen samples detected by the TCR-seq **b)** UMAP showing where clonotypes 302, 11155 and 21797 are residing. **c)** Volcano plot highlighting differentially expressed genes between liver and spleen detected for the clonotype 302 using pseudo bulk approach. log2FC positive means liver is upregulated relative to the control (spleen) **d)** UMAP showing where TRLs (from Fig.3 f,g, Fig.4) are residing.

## Discussion

UM continues to be a condition with unmet medical need, despite recent advances in locoregional therapy and immunotherapy^46^. However, the fact that a T cell engager is prolonging survival in patients with UM having an HLA-A2 genotype^14^ and that immune checkpoint inhibitors in various combination therapies on occasion can yield clinical benefit ^7,8,11^ suggests that if we learn more about the tumor immunology of UM, then we can devise new immunotherapies. Here, we generated datasets and models that can be used not only to characterize tumor immunity, but also to functionally test hypotheses and targets in autologous culture and *in vivo* systems.

ACT with TILs has also been tested in patients with metastatic UM^17^. Although there were responses in a few patients, the data clearly show room for improvement. The HAITILS trial (NCT04812470) addresses whether hepatic arterial infusion results in local accumulation of TILs in liver metastases. However, this will not circumvent the low mutational burden of UM (other than iris melanoma^22^) and the potentially low number of TRLs in the cell product (that we see in this study). A greater understanding of TRLs in patients’ metastases is therefore of interest to learn how to enhance the activity of TIL products and how to expand, for instance, neoantigen- or melanoma-associated antigen-specific TRLs^47,48^. Here, we aimed to identify tumor-reactive lymphocytes in terms of marker characteristics and functional assessments of TILs. These analyses suggest that some of the TILs in our cultures that became activated (4-1BB positive) when binding tumor cells were already TRLs (4-1BB or PD-1 positive) when they resided in the tumor. However, they were never able to clear the tumor, and most likely became exhausted or dysfunctional in the microenvironment.

Presence of TRL-specific markers adds validity to a strategy whereby enrichment of TRLs could be done directly from the tumor by sorting for those markers. Indeed, previous studies have employed activation, exhaustion, or tissue-resident markers such as 4-1BB^49^, PD-1^50–52^, CD39,^52,53^ and CD103,^54,55^ or the specificity of a TCR for a melanoma-associated or melanoma-specific antigen^56,57^. However, a risk with this strategy is that some TRLs are missed. Most notably, we revealed TRLs in clusters of T cells negative for activation and exhaustion markers. These cells are interesting since they express high levels of perforin and granzyme B, suggesting that they are cytotoxic. It is tempting to speculate that these cells are effector or effector memory cytotoxic T cells since they express RNA encoding CD16, FCRL6, and CX3CR1^58–60^. In the UM9 sample, most of the T cells in the biopsy that mapped to TRLs identified in the tumor sphere assay resided in this cluster.

PDX models have been developed for UM, but to our knowledge, this is the first time they have been used to study immunotherapy using autologous TILs and single-cell sequencing. Although one model responded to TIL therapy, the other model did not. In the UM22 liver metastasis model, we found evidence that T cells were pushed out of the tumor. This may partly explain the failure of immune checkpoint inhibitors in many patients ^61^ and why ACT with TILs does not work for the majority of patients with metastatic UM^17^. Nevertheless, we profiled TRLs that preferentially migrated to liver metastases in mice, as opposed to non-reactive T cells and those that migrate to the spleen. We identified marker genes and phenotypic states that might be used to identify relevant subsets present in patient biopsies and TIL cultures. These T cells appeared to resemble a recently described *IL7R*^-^*HAVCR2*^+^*KRT86*^+^*DUSP4*^+^*LAYN*^+^ subset, which has been associated with response to anti- PD1 therapy in lung cancer^45^. They also tended to express *ITGAE* (CD103), suggesting that they may be tissue-resident memory cells. This subset was identified in both PDX liver metastases, cultured TILs, and patient biopsies. The phenotypes of cells with this marker combination were predominantly exhaustion-like in the PDX samples as well as in biopsies. This is compatible the results from the profiling of biopsy CD8^+^ T cells, where mapping of experimentally identified MART1-reactive T cells back to biopsies and computational prediction of tumor antigen-recognizing T cells suggested an over-representation of TRL candidates in exhausted and late-activated gene expression-based clusters. Whether cells expressing these canonical exhaustion-related markers are truly dysfunctional or capable of eliciting an antitumor response remains to be explored^45^.

In the UM1 liver metastasis model, the tumor was eradicated, demonstrating that TILs can cause a complete response also of liver metastases. The single-cell data obtained confirmed the loss of melanoma cells but it also resulted in a loss of CD8+ T cells. It is well known in immunology that the fate of CD8 T cells after an acute immunological reaction is either death of the effector cells or development of memory. A strength of the PDX experiment is that we could model. But since we harvested the livers after tumor eradication, a limitation is that we could not investigate the expression of genes in T cells that are actively combating the tumor. However, since the CD4+ T cells were present in high numbers, we could compute differences between the T cells of the same clonotype residing either in liver or in spleen. This analysis did indeed suggest that some level of activation/exhaustion remained even after tumor clearance. Moreover, some of the TCRs from the coculture experiment of tumorspheres and TILs designed to reveal TRLs, were indeed identified in vivo among the CD8 and the proliferative cells. We are therefore confident that PDX models such as UM22 and UM1, will be useful in future studies where T cells are sorted for potential markers prior to expansion or infusion, where the T cells are genetically engineered or grown in cocktails of factors to enhance potency and stemness. We also propose the model to be appropriate for further humanization to unravel the interplay between tumor cells, liver cells and stroma cells, myeloid cells and other immune cells.

When mapping TCRs to TCR databases, we noted not only similarities to known melanoma-associated antigens, but also to viruses. Importantly, TCRs that recognize different antigens can share CDR3β chains. The TCR alpha chain is even more promiscuous but not always reported in these databases, which makes it difficult to find accurate TCR pair matches. Therefore, until current databases have been expanded and improved, we cannot be certain about the exact antigen matches of some of these receptors. Nevertheless, an increasing body of literature suggests that bystander cells in tumors are often antigen-experienced, can be directed against bacteria or viruses, and even participate in tumor control, for instance during immune checkpoint inhibitor therapy^62–64^. Whether the presence of foreign antigen-specific T cells is beneficial for cellular immunotherapy remains to be evaluated.

In summary, we profiled the tumor microenvironment of metastatic uveal melanoma and identified the phenotypes of tumor-reactive T cell subsets. The results of this study provide new avenues for targeted expansion of T cells to improve cellular and immune checkpoint immunotherapy. Importantly however, to potentiate TIL therapies in UM, immune cold tumors likely need to be turned hot, i.e., to create an inflammatory environment promoting T cell infiltration. Simultaneously, lymphodepletion is needed to avoid existing T cells sequestering IL2 needed for the TIL infusion product. Tumor-targeting treatments such as isolated or percutaneous hepatic perfusion (IHP and PHP) may fulfill these needs and could act as adjuvant therapy for both immune checkpoint inhibitor treatment and TIL therapy. As melphalan triggers immunogenic death, influx of new T cells into cold tumors may occur, partly explaining the very high response rate of these therapies^15^. Ongoing clinical trials are combining locoregional therapy with immune checkpoint inhibitors (NCT04463368) to capitalize on this concept. However, data from these trials have not yet been published. Combination of liver-directed therapy and TIL therapy is also warranted in future animal experiments and clinical trials. We propose that PDX models are optimal to capture the intra-patient heterogeneity of cancer and that multiple mechanisms in tumor immunology and treatment can be modelled at single-cell resolution.

## Methods

### Patient samples

According to the ethical approval of the Regional Ethical Review Board (#289-12, #44-18 and #144-13), the patients provided oral and written information with signed informed consent agreements. Biopsies were extracted from either subcutaneous or liver metastases. Tumor biopsies were divided into pieces that were snap-frozen, minced, embedded in formalin blocks, or used for tumor-infiltrating lymphocyte cultures and cryopreservation. Snap-frozen tumor fragments were homogenized using a Bullet Blender (Next Advance, NY, USA). DNA and RNA were extracted using the AllPrep DNA/RNA kit (Qiagen, Germany).

### Immunostainings of sections

Formalin-fixed, paraffin-embedded (FFPE) patient samples were evaluated using H&E staining and IHC. IHC was performed using an autostainer (Autostainer Link 48, Dako) with primary anti-human antibodies against Melan-A (clone A103; Dako), S100 (IR504; Dako), PRAME (E7I1B, CST), SOX10 (E6B6I, CST), HMB45 (M0634; Dako), CD3 (D4V8L, CST), and CD137 (E6Z7F, CST). HRP DAB or HRP Magenta (Dako) was used to stain the proteins of interest, and counterstaining was performed using hematoxylin.

We also performed multiplex immunofluorescence using The PhenoCycler-Fusion system (Akoya Biosciences) at the Spatial Proteomics Facility at SciLifeLab, Stockholm. The data for CD3, ICOS, and PD-1 primary antibodies are shown here; however, the full dataset will be published elsewhere.

### PDX models

Animal experiments were conducted in conformity with E.U. directive 2010/63 (Regional Animal Ethics Committee of Gothenburg approvals no. 2014-36 and 2018-1183). PDX models were generated by transplanting small pieces into the flank (subcutaneous) of immunocompromised, non-obese, severe combined immune deficient interleukin-2 chain receptor γ knockout mice (NOG mice) or hIL2-NOG mice (Taconic, Denmark). Cryopreserved single cells from biopsies were thawed and transplanted by splenic injection to create liver metastases in NOG mice or hIL2-NOG mice (Taconic, Denmark). All PDX tumors were analyzed using IHC, and their identity was verified by staining patient biopsies using clinically graded antibodies. For subcutaneous implants, tumor growth was monitored twice a week following TILS treatment. After establishment, the liver metastasis models formed liver metastases equally well by splenic injection and tail vein injection. The mice were evaluated using ultrasound scans before TIL treatment and grossly assessed at the time of sacrifice.

### Generation of TILs and MART1-specific T cells

The metastatic fragments were placed in culture medium (90% RPMI 1640 (Invitrogen), 10% heat inactivated Human AB serum (HS, Sigma-Aldrich), 6000 IU/ml recombinant human IL2 (Peprotech), penicillin and streptomycin (Invitrogen) to generate young TILS (yTILs) as previously described ^32^. MART1 specific yTILs were identified as previously described ^22^ and sorted using FACSAria III (BD Biosciences). yTILs were expanded using a standard REP (Rapid Expansion Protocol), constituting irradiated (40 Gy) allogeneic feeder cells, CD3 antibody (clone: OKT3) (30 ng/ml) (Miltenyi), medium (AIM-V, Invitrogen) supplemented with 10% HS and 6000 IU/ml IL2 and monitored for 14 days with media changes ^32^. After completion of the REP, MART1 specificity was confirmed using MART1-specific dextramers (Immudex, Copenhagen, Denmark). REP and REP MART1 TILs were used for downstream, cell line, spheroid, and *in vivo* experiments.

### Spheroid assay

PDX models were harvested, and tumor tissue was digested using a human tumor dissociation kit (Miltenyi) according to the manufacturer’s instructions in conjunction with the gentleMACS Octo tissue dissociator (Miltenyi). This single-cell suspension was used to prepare spheroids using an ultra-low attachment (ULA), 96- well, clear, round-bottomed plates (Corning,NY), and GBM-MG serum-free media (CLS cell lines service, Germany) for 72-96 hours in a 37°C incubator. To monitor treatment responses, cells were labeled with CellVue™ Jade Cell Labeling Kit (Thermo Fisher Scientific) and/or CellVue Claret Far Red Fluorescent Cell Linker Mini Kit (Sigma) and used for co-culture experiments. The 3D images were captured using IncuCyte® (Essen BioScience) +/- TILs treatment, followed by imaging and analysis at 72 h with IncuCyte® Live Cell Analysis. Tumor spheres stained with a far-red dye were measured for the red object area/µm^2^ mask with >100 μm. Flow cytometry was used to confirm the positively stained immune cells (Jade-FITC channel) and cancer cells (far red-APC channel).

### Co-culture and TCR sequencing

A human UM cell line derived from patient UM22 ^22^ was grown in complete medium (RPMI-1640 supplemented with 10% FBS, glutamine, and gentamycin) and cultured at 37°C with 5% CO2. The cell line or spheroids were analyzed with or without TILs, and the co-culture was analyzed using human CD3 (FITC and HIT3a, BioLegend), CD8 (APC and SK1, BioLegend), CD107a (APC and H4A3, BD Biosciences), CD137 (PE and 4B4-1, BioLegend), active caspase-3 (clone C92-605; BD Biosciences), and granzyme B (clone GB11; BD Biosciences) antibodies for 30 min at 4 °C. Flow cytometry data were acquired using the BD Accuri C6 and BD Accuri C6 Plus (BD Biosciences). The supernatants of these samples were subjected to Granzyme B secretion analysis using an ELISA kit (R&D Systems). Following detection of the activated cell populations in certain samples, a microfluidics-based flow cell sorting instrument (WOLF NanoCellect, San Diego, USA) was used to sort CD3^+^CD137^+^ cells from the co-culture of tumor spheres and TILs. The RNA of the sorted cells was extracted using a Nucleospin RNA XS kit (Macherey-Nagel), and its concentration was measured using QuBit. Libraries were prepared for TCR profiling using a SMARTer Human TCR a/b Profiling Kit v2 (Cat. No. 63478) and subsequent NGS library preparation and bead purification, Tapestation 4200 (Agilent) was used to confirm sample quality control with automated electrophoresis.

### Statistical analysis for spheroid assays

Statistical tests for the spheroid assays were done using two-tailed unpaired *t*-tests, with *p*-values represented as * for *p* < 0.05 and ** for *p* < 0.01; all error bars represent standard error (SEM), unless otherwise stated.

### Single-cell RNA-seq data analysis

#### Library preparation

5’ GEX and TCR sequencing: Single cell suspensions prepared from cryopreserved uveal melanoma biopsies were subjected to dead-cell removal using beads (Miltenyi Biotech) before quality and viability validation and loading on Chromium instrument (10x Genomics). Single cell 5’ gene expression and TCR libraries were prepared using vendor protocol with the Next GEM Single Cell 5’ Kit v2 (10x Genomics, catalog number 1000263), Library Construction Kit (Catalog number 1000190) Single Cell Human TCR Amplification Kit (Catalog number 1000252) and Dual Index kit TT Set A (Catalog number 1000215). Libraries were sequenced for quality check on iSeq100 (Illumina) before deep sequencing on NovaSeq 6000 (Illumina) in the required format to yield at least 20,000 reads per cell for Gene expression and 5,000 reads per cell for TCR libraries.

#### Alignment and gene expression quantification

Paired single-cell RNA- and TCR sequencing data from biopsies were aligned to the 10x Genomics-provided GRCh38 reference genome (v. 2020-A) and V(D)J Reference (v. 7.0.0) using the Cell Ranger pipeline (v. 7.0.1) and the “multi” command, with both “Gene Expression” and “VDJ-T” analyses activated, as well as with the setting “check-library-compatibility,false”.

#### Removal of ambient RNA

To remove ambient RNA, CellBender (v. 0.2.0)^65^ was used on the output from Cell Ranger, with the parameters “--fpr 0.01 --epocs 150” together with expected and total cell numbers estimated from Cell Ranger quality control plots, as described in the instructions at https://cellbender.readthedocs.io/en/latest/usage/index.html (accessed 15 Sept 2022). The values for the parameters “--low-count-threshold” and “--learning-rate” were changed from their defaults for some problematic samples and iteratively adapted as needed after inspecting CellBender output quality control plots, as described in the documentation.

#### Separation of human and mouse reads in PDX scRNA-seq data

To separate reads of mouse and human origin in the scRNA-seq data from PDX samples, XenoCell (v. 1.0)^66,67^ was used. As input, compressed fastq files, merged across lanes, were provided, together with reference genomes for human and mouse (the same genomes as for Cell Ranger). An index was created with the command “xenocell.py generate_index”, with the parameters “--threads 1 --memory 24”. Reads were classified according to host (mouse) or graft (human) with the command “xenocell.py classify_reads” and the parameters “-- barcode_start 1 --barcode_length 16 --threads 1 -memory 24 --compression_level 1”. Reads belonging to human and mouse, respectively were then extracted into new fastq files using the command “xenocell.py extract_cellular_barcodes” with the same values for barcode_start, barcode_length, threads, memory and compression_level parameters as in the previous step, but with “--lower_threshold 0 --upper_threshold 0.1” for graft and “--lower_threshold 0.9 --upper_threshold 1.0” for host. The workflow was implemented in a Nextflow script available at (https://github.com/jowkar/xenocell_nextflow). The new fastq files were then used for separate Cell Ranger runs for human and mouse reads for a given sample, as described above for human biopsy samples, except with the corresponding 10x Genomics-provided mouse reference genome instead for reads determined to be of mouse origin.

#### Quality control and cell filtering

Gene counts and assembled TCR sequences where imported to R with Seurat^68^. An initial object was created and normalized with the NormalizeData function of the same package. Samples were integrated with fastMNN^69^ (batchelor package v. 1.10.0) via the convenience function RunFastMNN from the SeuratWrappers (v. 0.3.0) package (parameters: “features = 4000, auto.merge = T, d = 150, k = 15”). Assembled and annotated TCR sequences from Cell Ranger were further added to this object using djvdj (v. 0.0.0.9000, https://github.com/rnabioco/djvdj). Cells with < 200 reads were then removed.

A first set of potential doublets were identified using scDblFinder (v. 1.8.0), using default parameters. In addition to this, cells expressing conflicting lineage-associated markers were also marked as doublets. Incompatible marker sets were defined as any among CD3D, CD4, CD8A, CD8B or with assembled TCR receptor expressed together with any of MLANA, PMEL, TYR, HBA1, HBA2 or HBB; any cell with assembled TCR receptor expressed together with any of CD14, CD19, MS4A1, or JCHAIN; or any among MLANA, PMEL or TYR expressed together with CD14, CD19, MS4A1, JCHAIN or NCR1. For the purpose of further refining doublet detection, an initial clustering was performed using the Seurat functions FindNeighbors (parameters: ‘reduction = “mnn”, dims = 1:150, k.param = 5’) and FindClusters (“resolution = 5”). For each cluster, a two-sided Fisher’s exact test was carried out, assessing whether that cluster was associated with doublets. p-values were adjusted for multiple testing with Benjamini-Hochberg correction. In addition, each cluster was also assessed for higher than expected UMI counts per cell. Any cluster with Fisher test odds ratio < 1 and q < 0.05, percent doublets greated or equal to the 95% percentile of doublet percentages, or UMI counts above the 95% percentile were further nominated as a doublet cluster to be removed. These thresholds were chosen in conjunction with visual inspection of plots regarding how these candidate clusters associated with other metrics of poor quality, such as proximity in UMAPs to cells expressing incompatible lineage markers and higher/lower than usual ribosomal or mitochondrial read percentage.

Global thresholds to keep cells fulfilling the following criteria were set: number of detected genes > 200, UMI counts > 500, ribosomal read percentage < 40% and percentage of genes in the top 100 expressed genes < 80%. However, further specific filtering was made for two cell types, erythrocytes and plasma cells, known to have generally lower RNA content. Since common global thresholds on RNA and number of detected genes may filter out these cell types entirely, lower thresholds were motivated specifically for these. To establish suitable thresholds, candidate members of these cell types were first identified based on the distinctly expressed marker genes HLA1 (erythrocytes) and JCHAIN (plasma cells, with IL3RA and LILRA4 additionally demanded to be zero), respectively. Cells in each respective group (without having removed candidate doublets) were clustered based on Spearman correlation coefficients with the pheatmap function in the R package of the same name (default parameters) and visualised together. In both cases, cells were divided into clear clusters, some of which had predicted doublets were overrepresented. The latter clusters also tended to have higher than expected RNA content. Cluster without overrepresented candidate doublets were retained and the rest discarded from the dataset. Thresholds to keep cells with UMI counts and number of detected genes greater than the first quartile were defined for each of the two cell types based on cells in clusters not associating with doublets.

Further to this, additional quality filtering thresholds for remaining cells were determined adaptively using miQC (v. 1.2.0)^70^, which first creates mixture models to assess the distribution of number of genes detected versus percent mitochondrial reads in each sample and then filters cells based on a posterior probability of being compromised derived from this distribution. For a few samples that failed to converge with the default model, a one-dimensional instead of mixture model was instead used. For all samples, the parameters “posterior_cutoff = 0.9” and “keep_all_below_boundary = TRUE” were used to filter out low-quality cells.

Sample integration, unsupervised clustering and cell type annotation

After removing low-quality cells and doublets, all samples were reintegrated using fastMNN, with the same parameters as described above. A UMAP embedding was created using the Seurat function RunUMAP (parameters: reduction = “mnn”, dims = 1:150). Unsupervised clustering was performed with FindNeighbors (reduction = “mnn”, dims=1:150) and FindClusters (resolution = 2). Genes differing between clusters were assessed using the FindAllMarkers function (default parameters). Clusters were annotated based on a combination of generally known cell type markers (CD3D and CD3G for T cells, CD8A for CD8^+^ T cells, CD4 for CD4^+^ T cells, FOXP3 for regulatory T cells, NCAM1 and NCR1 for NK cells, CD19 and MS4A1 for B cells, HBA1, HBA2 and HBB for erythrocytes, CD14 for general monocytic cells, PMEL, MLANA, and TYR for melanocytic cells, as well as MKI67 for actively cycling cells) and markers compiled from literature^28,71–74^. CD8^+^ T cells where determined and studied in greater detail to identify further subgroups. After subsetting the main Seurat object to only contain CD8^+^ T cells, these were subclustered by rerunning FindNeighbors (‘reduction = “mnn”, k.param = 15, dims=1:150’) and FindClusters (“algorithm = 1, resolution = 1.3”). A new UMAP was created using RunUMAP (‘reduction = “mnn”, dims=1:150, min.dist = 0.05, n.neighbors = 15’). 16 clusters were obtained, among which some were merged based on similar expression of key marker genes. The new CD8^+^ T cell clusters were annotated based on a combination of literature-derived markers^28,75–77^ for CD8^+^ T cell phenotypes as well as genes found to distinguish the clusters that could be related to specific phenotypes.

#### Processing of scRNA-seq from TIL cultures

Single-cell RNA-seq data of TIL cultures from a previous study^41^ were obtained, loaded into a Seurat object in R and normalized with the NormalizeData function (default parameters). Variable features were determined using FindVariableFeatures (selection.method = “vst”, nfeatures = 4000). Cell cycle scores were then calculated using CellCycleScoring (default provided gene sets) and data scaled using ScaleData (vars.to.regress = c(“S.Score”, “G2M.Score”)). Prinicipal components were calculated using the RunPCA function (npcs = 30) and a UMAP embedding calculated using the RunUMAP function (dims = 1:30). These steps were performed on each sample separately, without sample integration.

#### GLIPH2 analysis

TCR sequences were clustered into inferred specificity groups using GLIPH2^43^. As input, unique TCR sequences and their counts from annotated CD8^+^ T cells were given, together with previously published HLA genotypes from bulk RNA and WGS^41^. HLA genotypes for two additional samples were inferred from previously published exome data^78^ using nf-core/hlatyping (v2.0.0)^79^, with the parameter “--genome GRCh37”. Clonotypes with more than one TCR alpha chain were excluded from analysis, as these are not supported by GLIPH2. Parameters provided were the option to use the v2 human reference for CD8 T cells and with “all_aa_interchangeable” set to Yes. The tool was run from the web interface and according to instructions at http://50.255.35.37:8080/ (accessed 21 Dec 2022). High-confidence TCR clusters were considered those with V-gene bias score < 0.05, at least 3 unique TCRs and at least 3 unique individuals contributing TCRs.

#### Copy number analysis

To determine copy number changes in UM cells from scRNA-seq data, the inferCNV R package (v. 1.10.1)^80^ was used according to instructions provided at https://github.com/broadinstitute/infercnv/wiki (accessed 2 Nov 2022). A genomic position file was created from the reference annotation provided with Cell Ranger (v. 2020-A) using the inferCNV helper script gtf_to_position_file.py. All cells annotated as UM were marked as “malignant” and remaining cells were used as a reference diploid background as input to inferCNV, together with the following parameters: ’cutoff=0.1, cluster_by_groups=TRUE, denoise=TRUE, HMM=FALSE, num_threads = 1, analysis_mode=“samples”, output_format = “pdf”’.

#### Differential expression analysis

Differentially expressed genes between experimental conditions in scRNA-seq data from PDX samples were assessed using a pseudo-bulk approach together. First, the all human CD8+ T cells for each sample were used to create gene subset expression matrix. All cells in a given sample were summed on a per-gene basis to create pseudo-bulk samples, imitating a bulk RNA-seq experiment. These samples were normalized and analysed for differential expression between the conditions represented by the samples using DESeq2^81^, with the parameters “test = “LRT”, reduced = ∼ 1”, relative to a full model specification encoding the two experimental conditions to be compared. For each pairwise comparison of conditions, only samples in those two conditions were included in the DESeq2 main object. To extract a table of results, the “results” function from the same R package was used, with the parameter “alpha = 0.05”. Significant genes were visualized with the pheatmap function from the R package of the same name (v. 1.0.12), with the parameter ’scale = “row”’.

#### scRNA and TCR-seq analysis for UM1 sample

FASTQs files lane-1 and lane-2 were concatenated and used as an input for the XenoCell (v.1.0) tool to split scRNA-seq data on host (mm) and graft (hs). XenoCell classify_reads function was used to calculate the percentage of graft- and host-specific reads for each cellular barcode. Then, extract_cellular_barcodes function was used to extract cellular barcodes for the host (mm), and the same function was used to extract cellular barcodes for the graft (hs). The feature-barcode matrix for graft (hs) was generated using cellranger multi (v.7.0.1) pipeline. Parameters --localcores and --localmem were set to 32 and 240 respectfully and human reference genome GRCh38 (v.2020-A) and V(D)J reference (v.7.0.0) were used. Next, cellbender remove- background CellBender (v.0.2.0) function was used to remove ambient RNA. For each sample cellbender parameters were adjusted as follows --expected-cells (25L: 7573; 25S: 2274; 39L: 7973; 39S: 10527; 45L: 8438; 45S: 10447; 55L: 6318; 55S: 4143), --total-droplets-included (25L: 17000; 25S: 10000; 39L: 12000; 39S: 17000; 45L: 16000; 45S: 19000; 55L: 15000; 55S: 13000), --fpr (25L: 0.01; 25S: 0.01; 39L: 0.01; 39S: 0.01; 45L: 0.01; 45S: 0.01; 55L: 0.01; 55S: 0.01), --epochs (25L: 150; 25S: 150; 39L: 200; 39S: 150; 45L: 150; 45S: 200; 55L: 150; 55S: 150), --low-count-threshold (25L: 3; 25S: 5; 39L: 3; 39S: 3; 45L: 3; 45S: 5; 55L: 5; 55S: 5), --learning-rate (25L: 0.0000125; 25S: 0.00005; 39L: 0.00000625; 39S: 0.0000125; 45L: 0.00000625; 45S: 0.00000625; 55L: 0.00005; 55S: 0.00005).

R (v.4.3.2) was used for downstream analysis. Data generated by CellBender was parsed using Read_CellBender_h5_Multi_Directory function from scCustomize (v.2.0.1) package. Seurat object was created using CreateSeuratObject function from Seurat (v. 5.0.1) package with parameters min.cells and min.features set to 3 and 200 respectfully. This resulted in the Seurat object with 23,884 genes across 64,091 cells. V(D)J data was added to the Seurat object using import_vdj function from djvdj (v. 0.1.0) library. Doublets were removed using scDblFinder function from scDblFinder (v. 1.16.0) package that resulted in 57,200 cells. Cells with a low coverage were removed before using scDblFinder (nCount <200, https://bioconductor.org/packages/release/bioc/vignettes/scDblFinder/inst/doc/scDblFinder.html). QCs filtering was performed to remove dying/dead, any remaining empty droplets or multiples cells. Cells with a high gene deletion number could indicate doublets or multiplets, thus, cells that have unique gene counts over 6,500 or less than 400 were removed. Cells with a total read count (or UMI) less than 500 and more than 25,000 were filtered out. Finally, cells with mitochondrial counts more than 20% and ribosomal counts more than 15% were removed. Next, the data was normalized, scaled, and dimensionality reduction applied using NormalizeData, FindVariableFeatures (nfeatures = 2000), ScaleData, RunPCA. This step followed standardized Seurat workflow (“Data analysis workflow Integrative analysis in Seurat v5”, https://satijalab.org/seurat/articles/seurat5_integration, accessed 2024). Integration was performed using IntegrateLayers function from Seurat (v. 5.0.1) package with parameters set to method = CCAIntegration, orig.reduction = “pca”, new.reduction = “integrated.cca”. FindNeighbors (reduction = “integrated.cca”, dims = 1:70) and FindClusters (resolution =c(0.1 to 2.0) , cluster.name = “cca_clusters”) functions from Seurat (v. 5.0.1) package were used and resolution 0.6 (15 clusters) was selected. FindAllMarkers (logfc.threshold = 0.3, min.pct = 0.3, only.pos = TRUE, test.use = “wilcox”) function from Seurat (v. 5.0.1) package was used to find differentially expressed genes. To visualize results DimPlot, VlnPlot, FeaturePlot and DotPlot Seurat (v. 5.0.1) functions were used. For plotting the volcano plot we used EnhancedVolcano (v.1.20.0; accessible from https://github.com/kevinblighe/EnhancedVolcano), for plotting a bar plot ggplot2 (v.3.5.0) was used. To pseudobulk, AggregateExpression function from Seurat (v. 5.0.1) package was used. The gene list was compiled using marker genes coming from differential expression analysis and from the literature. Results were visualized as a heatmap using plotGroupedHeatmap function from the scater (1.30.1) package. To find clonotype abundance calc_frequency function from djvdj (v. 0.1.0) package was used, to visualize results plot_clone_frequency function was used from the same package.

### TCR sequence analysis

#### Bulk TCR sequencing data analysis

TCR sequences sequenced with the Takara SMARTer Human TCR a/b Profiling Kit v2 were aligned and assembled using the open source Cogent NGS Immune Profiler software (v1.0) provided by the same company, using the parameters “-r TCR” and “-t Both”.

#### Matching of TCR receptors between samples

TCRs from single-cell RNA-seq samples and/or bulk TCR-seq samples were matched based on identical combinations of CDR3β, Vb and Jb. This was the minimal common combination of criteria available on which to consistently compare TCRs across all TCR-sequenced samples, since bulk the TCR-seq protocol does not provide information in pairing of alpha and beta chains.

#### Matching of TCR receptors to public databases

To match TCRs to public databases with information on possible antigens they might recognize, TCRMatch (v. 1.0.2)^42^ was used (parameter: -s 0.97) together with a modified version of the reference database from IEDB^82^ provided with the tool, which we expanded with additional information from VDJdb^83^, McPAS-TCR^84^ and TCR3d^85^ (using the datasets for both cancer and virus-targeting TCRs). In addition, this database file was further modified to replace internal commas in antigen names, since the default formatting conflicts with the use of comma as the main delimiter in the database file. This facilitated easier analysis of the results. TCR matches were visualized with Sankey diagrams using the ggsankey R package (v. 0.0.99999, https://github.com/davidsjoberg/ggsankey).

#### Overrepresentation analysis for TCRs in CD8^+^ T cell clusters

To assess whether the 41BB- or MART1-reactivity experiment would enrich for clones mapping to any phenotype relative to cultured TILs in general, a binomial test was used. The test considers the observed fraction of cells with matches to the reactivity screen versus an expected probability. Given that cells from the reactivity experiment were taken from a TIL culture, we wanted to assess whether these had a meaningfully different distribution compared to cultured TILs in general in regards to cluster representation. Therefore, the expected probability was estimated as the fraction of TILs that match any given cluster. Thus, these two frequencies were compared with one two-sided binomial test (binom.test in R) per cluster, and *p*-values further adjusted for multiple testing with Bonferroni correction.

### Data access statement

The single-cell sequencing data has been uploaded to European Genome-Phenome Archive and is available under controlled access after publication. All remaining data are within the paper and its Supporting Information files or available on request.

## Author contributions

JK: conducted all bioinformatic analysis except for the UM1 liver metastasis model data, generated figures and wrote several sections of the paper; VS: conducted experiments, generated figures and wrote some sections of the paper; ROB: coordinated patient consents, surgeries and collection of liver metastases of uveal melanoma; MI: conducted single-cell sequencing; SA: performed cell sorting and flow analyses; SS: generated PDX models and developed new transplantation and imaging techniques; IK: conducted bioinformatic analysis for the liver metastatic model of the UM1 sample; AS: managed single-cell sequencing workflow; LN: supervised the study, coordinated patient consents and sample collection; LMN: supervised, managed the biobank and sequencing, generated TILs, managed, conceived and visualized data from animal experiments; JAN: conceived and supervised the study, wrote the paper and generated figures; all authors, read, edited and approved the manuscript.

## Acknowledgments

We thank Carina Karlsson for providing technical support. Grant support came from Cancerfonden (to J.A.N. and R.O.B), Familjen Erling Persson (to J.A.N.), Knut and Alice Wallenberg Foundation (to J.A.N. and R.O.B), Vetenskapsrådet (to J.A.N. and R.O.B.), Sjöbergstiftelsen (to J.A.N.), BioCARE Strategic grants (to J.A.N.), Lion’s Cancerfond Väst (to S.A. and L.N), Västra Götaland Regionen ALF grant (to J.A.N., R.O.B. and L.N.), Assar Gabrielsson fond (to V.S.), Gustaf V Jubileumsklinikens forskningsfond (to L.N.) and Wilhelm & Martina Lundgrens Vetenskapsfond (to J.K.). J.A.N. is the Inaugural Chair of Melanoma Discovery, primarily supported by donations to Perkins from family and friends of Scott Kirkbride, the MACA Cancer 200 ride for cancer research, the HBF Foundation and Perpetual. The Genomics WA facility (A.S.) is supported by BioPlatforms Australia, State Government Western Australia, Australian Cancer Research Foundation, Cancer Research Trust, Harry Perkins Institute of Medical Research, Telethon Kids Institute and the University of Western Australia. We also thank the Pawsey Supercomputing Centre for providing computational resources for this project.

## Supplementary Figure and Table Legends

**Supplementary Figure 1.**
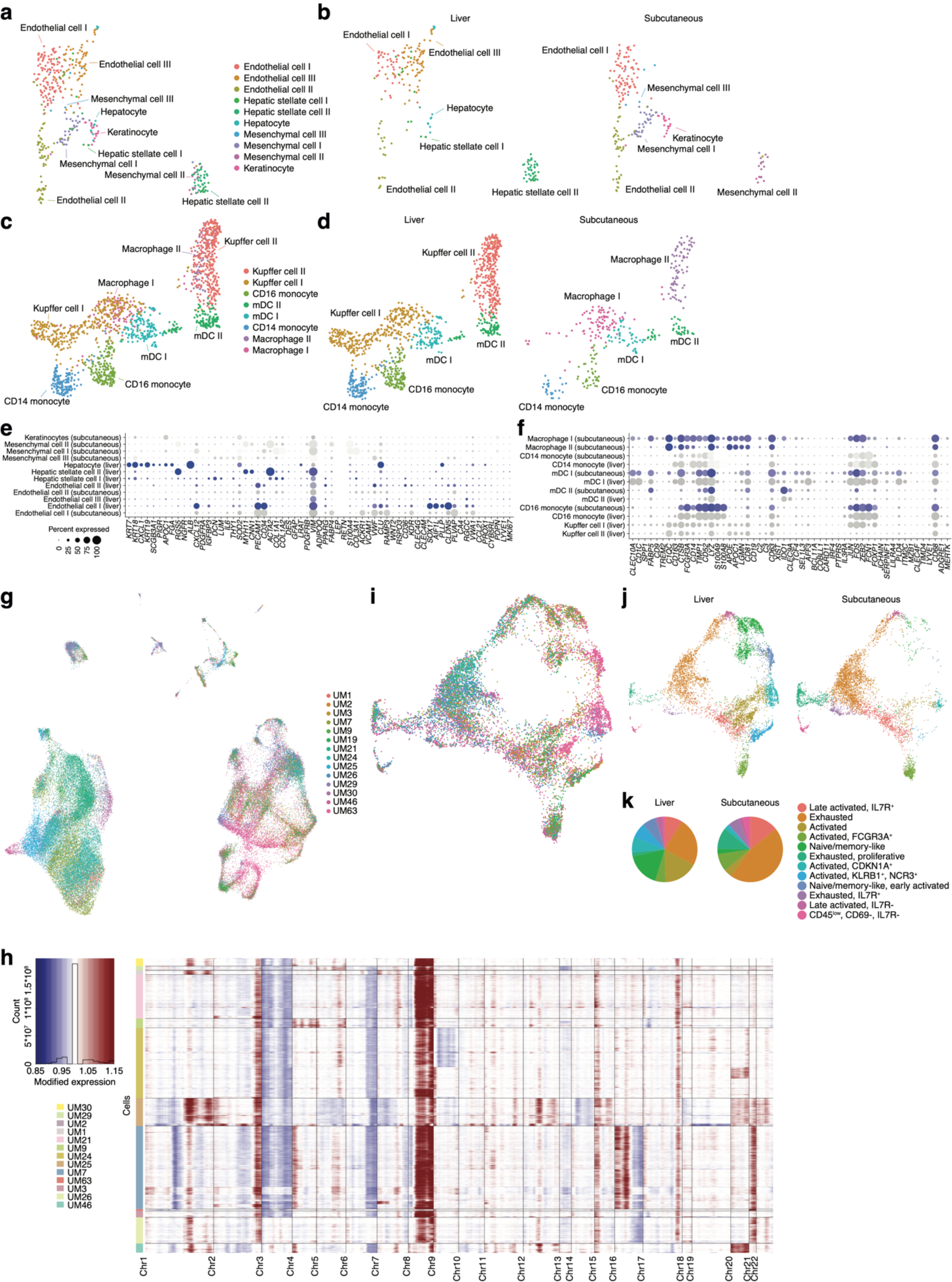
**a-b)** Subclustering of cells that initially grouped together with endothelial cells (a). Shown per tissue of origin (b). **c-d)** Subclustering of monocyte-like cells. Shown per tissue of origin (d). **e-f)** Marker genes used to label clusters in (a) and (c), respectively. **g)** Sample contribution to each cluster from Fig. 1b. **h)** Copy number profiles for UM cells identified in each sample, as determined by inferCNV^80^ analysis. Blue represents copy number loss and red copy number gain. **i)** Sample contribution to each cluster from Fig. 2a. **j-k)** Cell contribution to each CD8^+^ T cell cluster from biopsies of liver and subcutaneous tumors, respectively.^69^

**Supplementary Figure 2.**
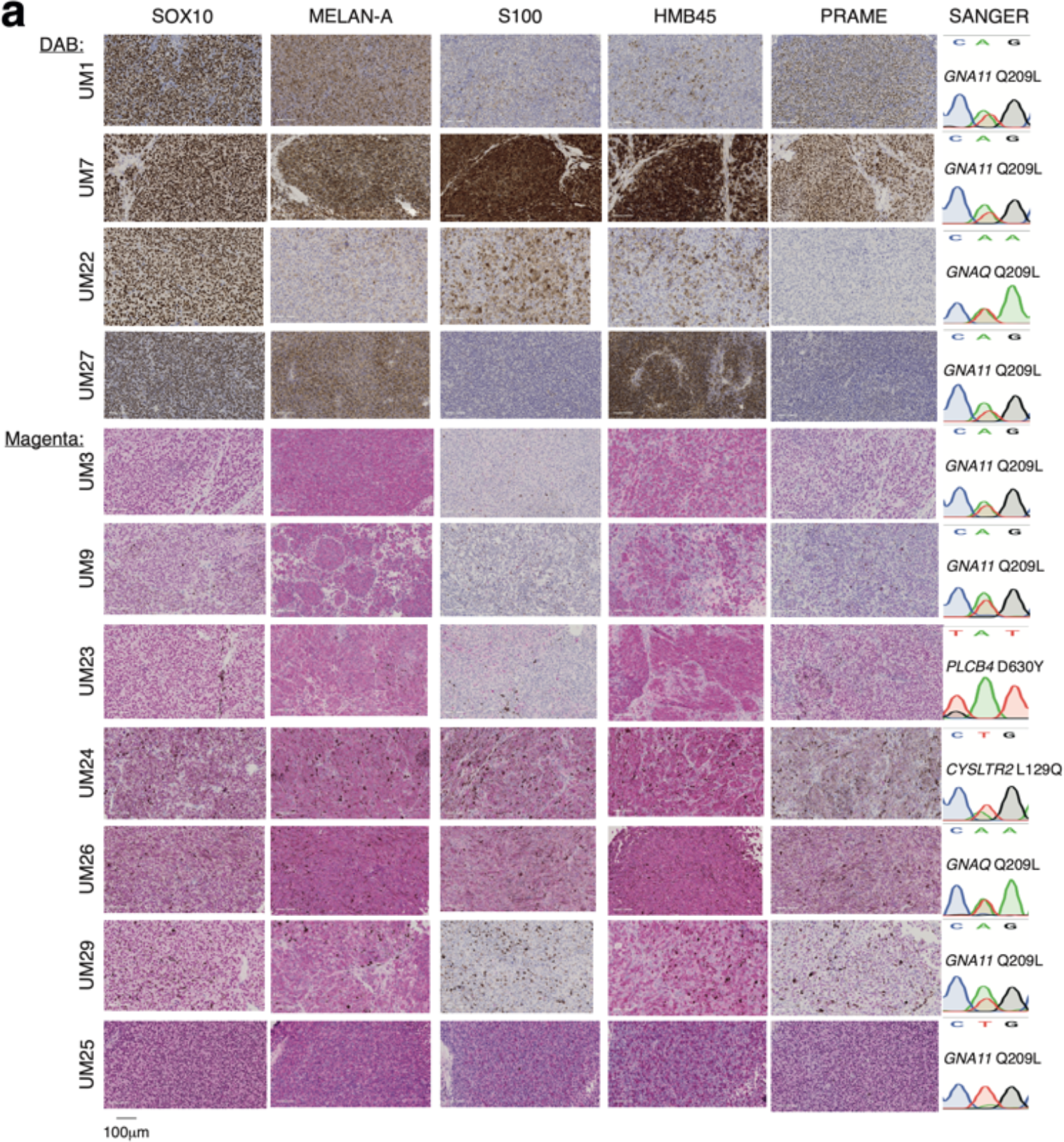
IHC showing expression of HMB45, SOX10, S100, MELAN-A, PRAME expression in PDX models. Staining color used was either DAB (top four samples) or magenta.

**Supplementary Figure 3.**
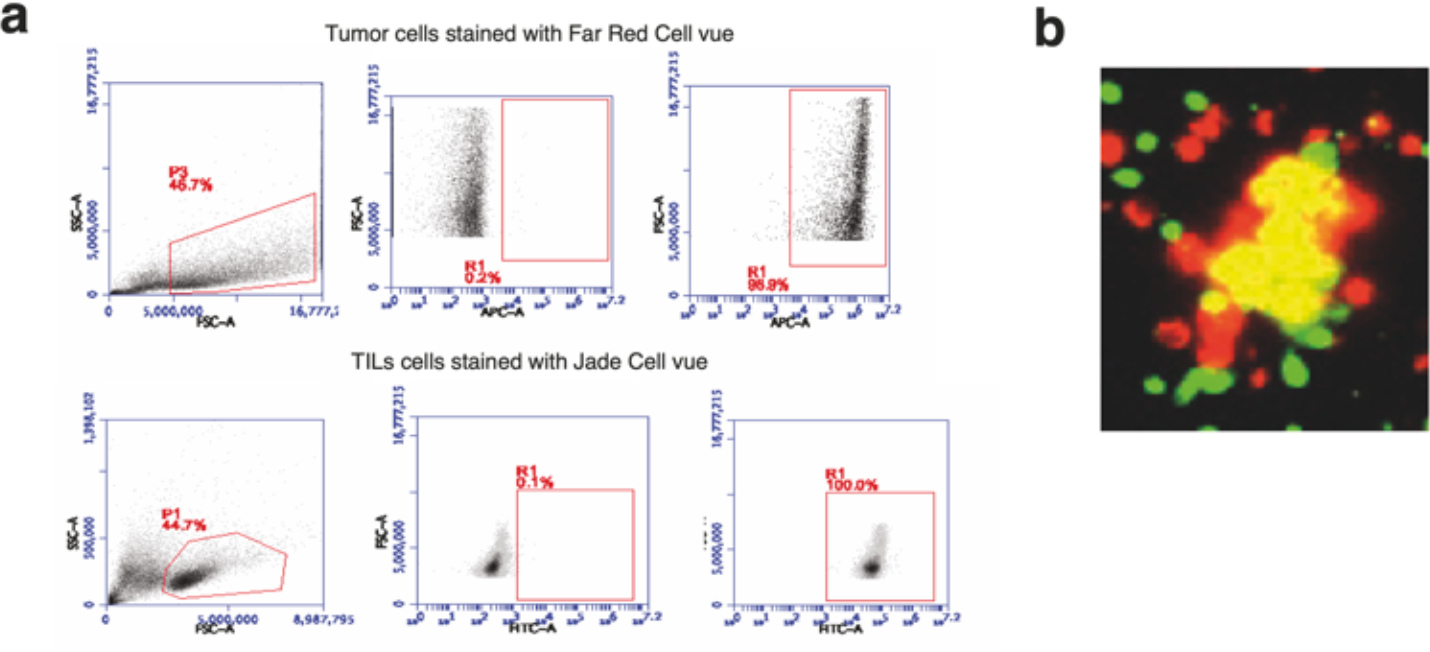
**a)** Gating strategy for Far Red Dye tagged tumor cells (APC) and Jade dye tagged TILs (FITC). **b)** Far Red fluorescent dye incorporated spheres co-cultured with Jade dye labeled TILs (green) co-culture with a co-localized population (yellow) at 24-hour time-point.

**Supplementary Figure 4.**
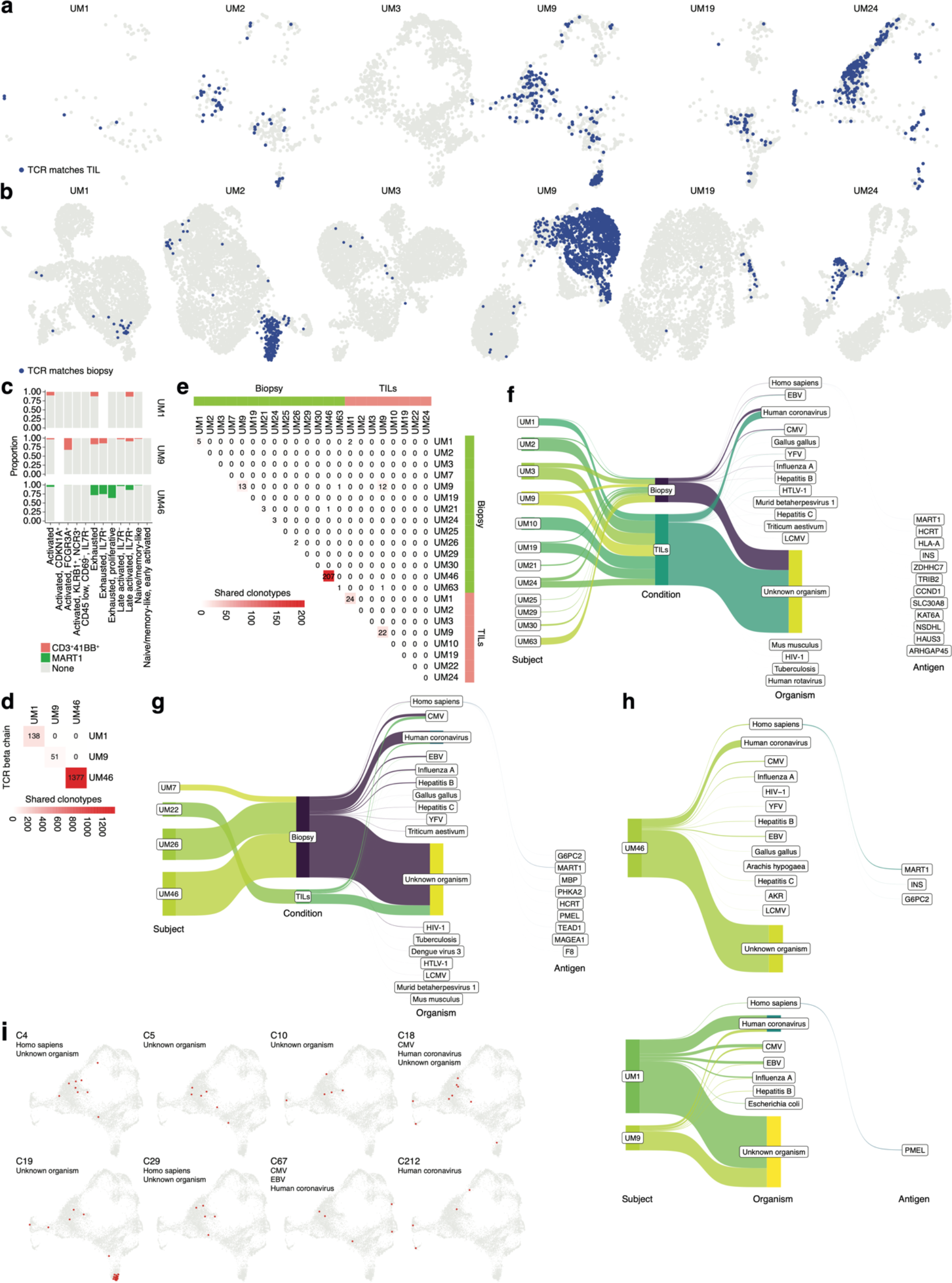
**a)** Biopsy CD8^+^ T cells with clonotypes that match scRNA-sequenced TILs, as in Fig. 4c but shown separately for each sample. **b)** TIL cells with clonotypes that match any corresponding biopsy. **c)** Proportions within each CD8^+^ T cell cluster that match either 4-1BB^+^ or MART1-selected TILs from UM1, UM9 and UM46, respectively. **d)** Overlap among TCRβ chains from the reactive cells sorted out from these three samples. **e)** Shared TCRβ chains among the subset of cells from biopsies and TIL cultures that also have a match to any TCRβ chain among experimentally identified reactive TILs. **f-g)** Matches to any CDR3β-antigen pair in public databases, as inferred by TRCMatch, for CD8^+^ T cells from biopsies and TILs. Samples from liver and subcutaneous metastases shown in (f) and (g), respectively. **h)** As in (f-g), for experimentally identified reactive TILs. **i)** Cells from biopsy CD8^+^ T cells that were included in any high-confidence GLIPH2 cluster.

**Supplementary Figure 5.**
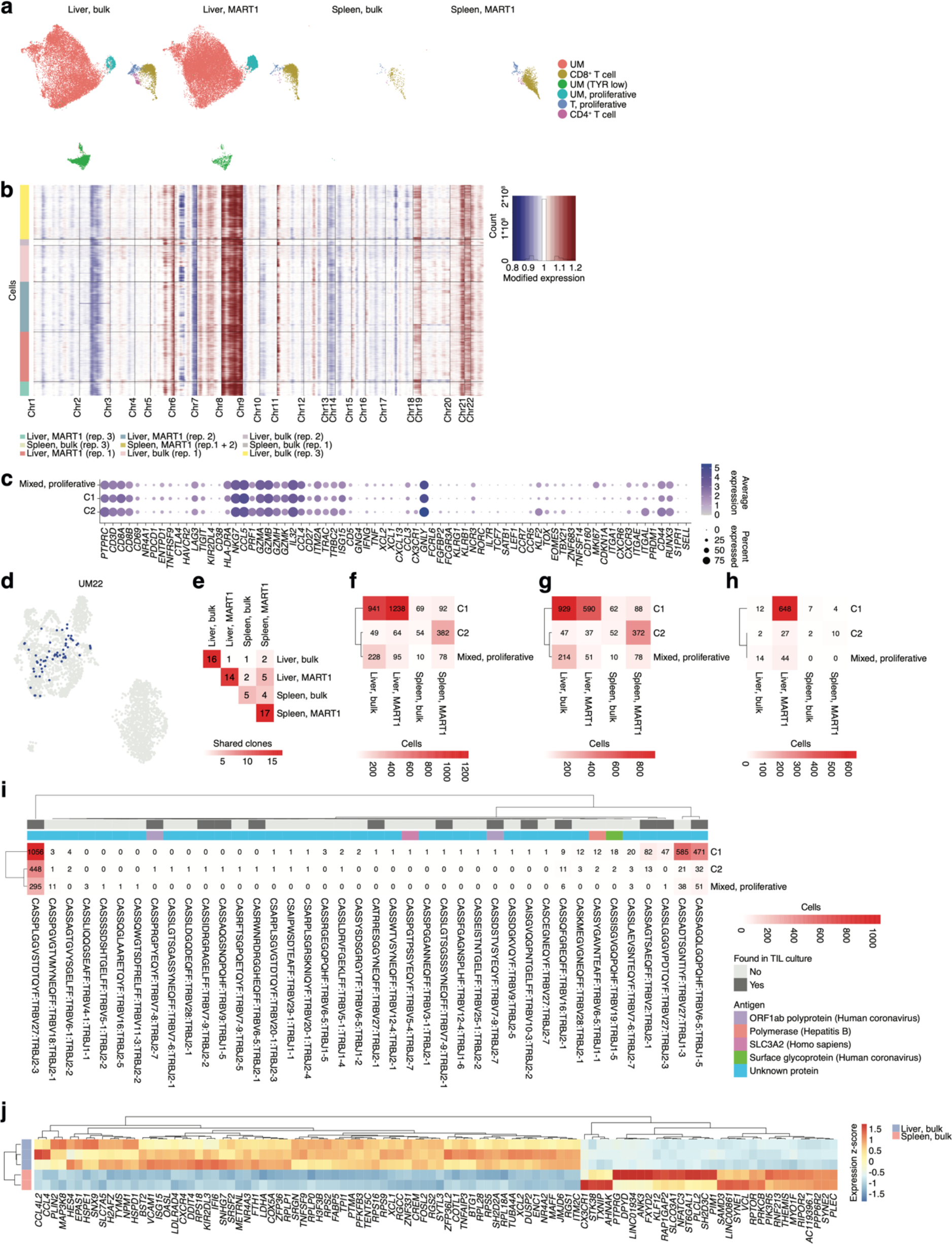
**a)** All cells from scRNA-seq analysis of PDX samples, shown separately per experimental condition. **b)** Copy number profiles of UM cells in PDX samples, as determined by inferCNV^80^ analysis. Blue represents copy number loss and red copy number gain. **c)** Markers discriminating CD8^+^ T cell clusters in biopsies, contrasted with CD8^+^ T cell clusters identified in PDX samples. **d)** TCRβ chains identified in the PDX samples that match clonotypes in a separate scRNA-seq experiment of UM22 TILs. **e)** Shared unique TCRβ chains between PDX samples from each experimental condition. **f)** As in (e), shown relative to PDX CD8^+^ T cell clusters. **g)** As in (f), but only including cells that also match any clonotype found in scRNA-seq from UM22 TIL cultures. **h)** Arithmetic difference between (g) and (f), showing cells with TCRs uniquely detected PDX samples compared to the sequenced UM22 TIL culture. **i)** All TCRβ chains identified in each PDX CD8^+^ T cell cluster. Clonotypes found in TIL cultures are highlighted, as well as any matches to public antigen databases. **j)** Differentially expressed genes between bulk TIL mixtures residing in liver and spleen of PDX models. A pseudo-bulk approach^86^ was used, summing read counts across all cells within a given replicate, and statistical testing performed with DESeq2^81^. Genes with *q* < 0.05 were considered significant.

**Supplementary Figure 6.**
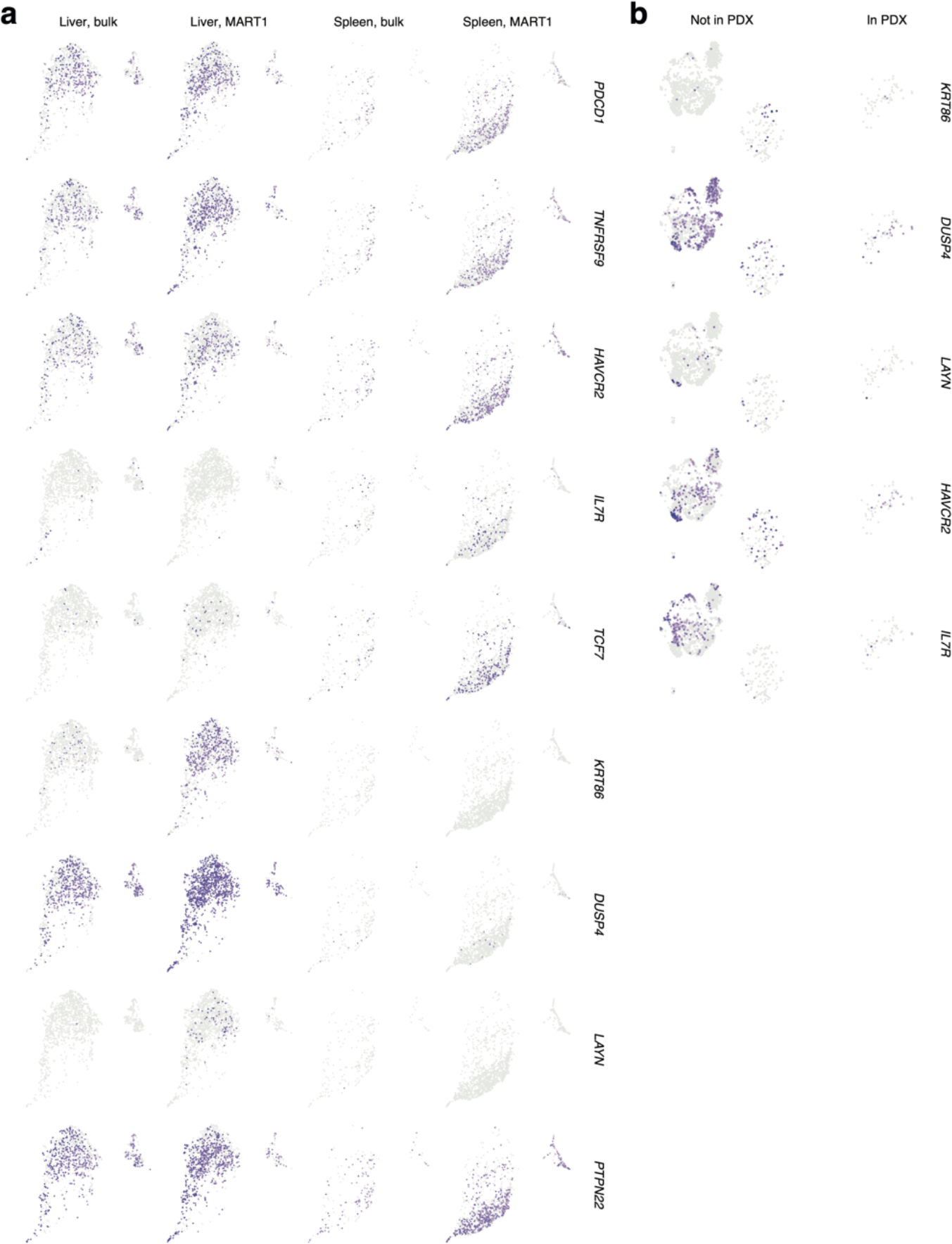
**a)** Expression of marker genes related to a previously described tumor-reactive set of tissue-resident CD8^+^ T cells^45^, among CD8^+^ T cells found in PDX models. **b)** Expression of *KRT86*, *DUSP4*, *LAYN*, *HAVCR2* and *IL7R* in TIL culture from UM22. Whether a given cell has a clonotype identified in PDX samples is indicated.

**Supplementary Figure 7.**
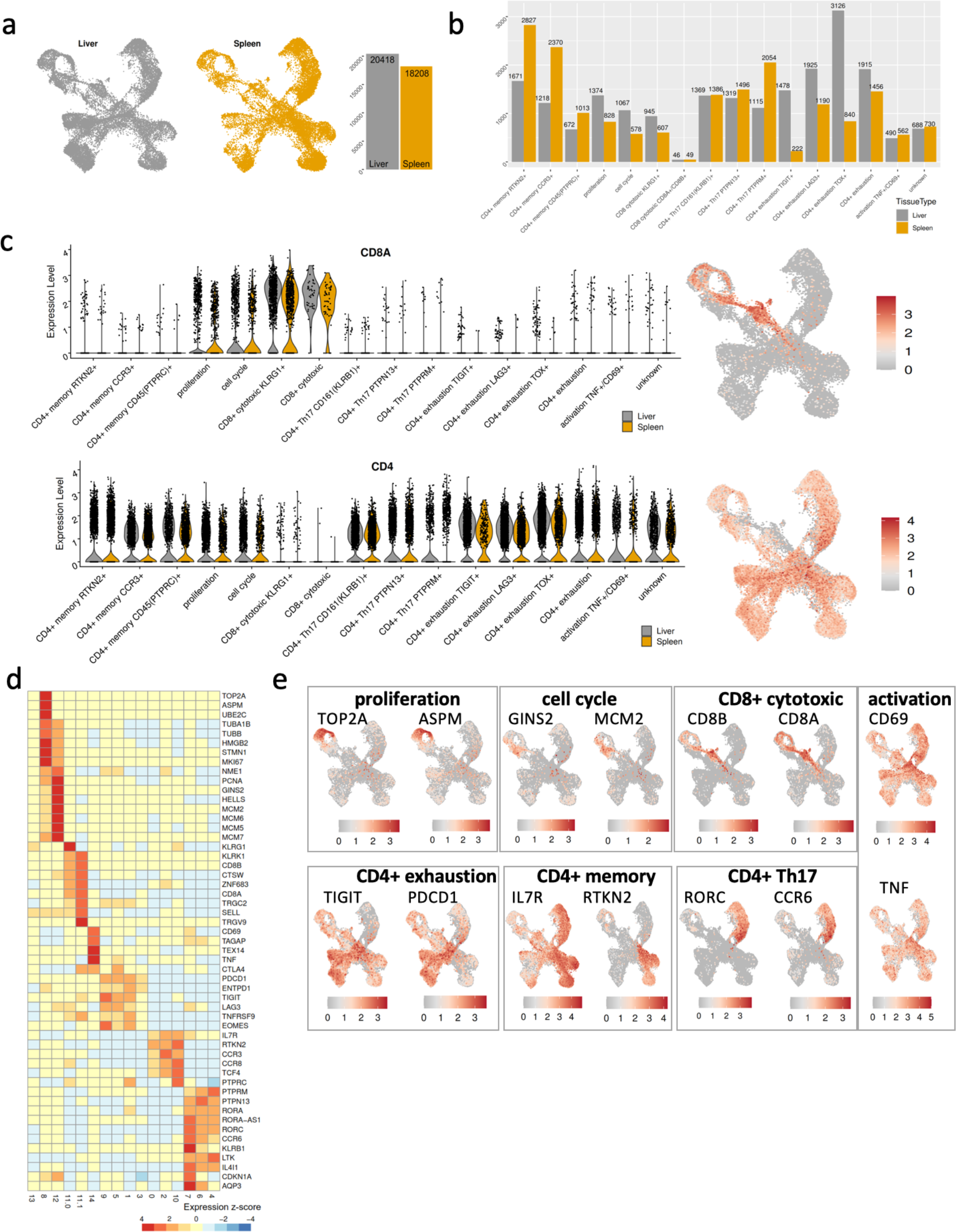
**a)** UMAP split on spleen and liver. Liver has 20,418 cells and spleen has 18,208 cells. **b)** Bar plot showing number of cells across each cell type comparing between spleen and liver. **c)** Violin plot showing expression distribution for CD4 and CD8A genes across clusters between liver and spleen. In liver, CD8A gene is mainly expressed in CD8+ cytotoxic KLRG1+ and CD8+ cytotoxic clusters, and partially in proliferative and cell cycle clusters. CD4 gene is expressed in all cell types except for cluster CD8 cytotoxic KLRG1+ and CD8 cytotoxic clusters. Two UMAPs showing expression level of CD4 and CD8A genes. **d)** Heatmap showing gene expression (z-score) values of pseudo bulk (“summed counts”) for each cell population. **e)** Feature plot showing gene expression of two common marker genes for each cell population.

**Supplementary Table 1.** Statistically identified marker genes for each CD8^+^ T cell cluster in Fig. 2a., using the FindMarkers function in the Seurat R package.

**Supplementary Table 2.** Complete statistics for the analysis of cluster representation among reactivity-screened TCRs matched to biopsy CD8^+^ T cells, as compared to representation among TILs matched to the same clusters (Fig. 4d).

**Supplementary Table 3.** Complete statistical output from the GLIPH2 analysis referred to in Fig. 4h**-i** and Supplementary Fig. 4i.

**Supplementary Table 4.** Complete statistics for the differential expression analyses in Fig. 5k and Supplementary Fig. 5j.

**Supplementary Table 5.** Differentially expressed genes in different clusters of the UMAP in Fig. 5.

**Supplementary Table 6.** References for marker gene selection and annotation of clusters in in Fig. 5.

## References

1. Jager, M. J. et al. Uveal melanoma. Nat Rev Dis Primers 6, 24 (2020).

2. Buder, K., Gesierich, A., Gelbrich, G. & Goebeler, M. Systemic treatment of metastatic uveal melanoma: review of literature and future perspectives. Cancer Med 2, 674–86 (2013).

3. Khoja, L. et al. Meta-analysis in metastatic uveal melanoma to determine progression free and overall survival benchmarks: an international rare cancers initiative (IRCI) ocular melanoma study. Ann Oncol 30, 1370–1380 (2019).

4. Carvajal, R. D. et al. Metastatic disease from uveal melanoma: treatment options and future prospects. Br J Ophthalmol 101, 38–44 (2017).

5. Johnson, D. B. et al. Response to Anti-PD-1 in Uveal Melanoma Without High-Volume Liver Metastasis. J Natl Compr Canc Netw 17, 114–117 (2019).

6. Algazi, A. P. et al. Clinical outcomes in metastatic uveal melanoma treated with PD-1 and PD-L1 antibodies. Cancer 122, 3344–3353 (2016).

7. Piulats, J. M. et al. Nivolumab Plus Ipilimumab for Treatment-Naive Metastatic Uveal Melanoma: An Open-Label, Multicenter, Phase II Trial by the Spanish Multidisciplinary Melanoma Group (GEM-1402). J Clin Oncol JCO2000550 (2021) doi:10.1200/JCO.20.00550.

8. Pelster, M. S. et al. Nivolumab and Ipilimumab in Metastatic Uveal Melanoma: Results From a Single-Arm Phase II Study. J Clin Oncol JCO2000605 (2020) doi:10.1200/JCO.20.00605.

9. Najjar, Y. G. et al. Ipilimumab plus nivolumab for patients with metastatic uveal melanoma: a multicenter, retrospective study. J Immunother Cancer 8, (2020).

10. Jespersen, H. et al. Concomitant use of pembrolizumab and entinostat in adult patients with metastatic uveal melanoma (PEMDAC study): protocol for a multicenter phase II open label study. BMC Cancer 19, 415 (2019).

11. Ny, L. et al. The PEMDAC phase 2 study of pembrolizumab and entinostat in patients with metastatic uveal melanoma. Nat Commun 12, 5155 (2021).

12. Damato, B. E., Dukes, J., Goodall, H. & Carvajal, R. D. Tebentafusp: T Cell Redirection for the Treatment of Metastatic Uveal Melanoma. Cancers (Basel) 11, (2019).

13. Middleton, M. R. et al. Tebentafusp, a TCR/anti-CD3 bispecific fusion protein targeting gp100, potently activated anti-tumor immune responses in patients with metastatic melanoma. Clin Cancer Res (2020) doi:10.1158/1078-0432.CCR-20-1247.

14. Nathan, P. et al. Overall Survival Benefit with Tebentafusp in Metastatic Uveal Melanoma. N Engl J Med 385, 1196–1206 (2021).

15. Olofsson Bagge, R., et al. Isolated Hepatic Perfusion With Melphalan for Patients With Isolated Uveal Melanoma Liver Metastases: A Multicenter, Randomized, Open-Label, Phase III Trial (the SCANDIUM Trial). J Clin Oncol JCO2201705 (2023) doi:10.1200/JCO.22.01705.

16. Ben-Shabat, I. et al. Long-Term Follow-Up Evaluation of 68 Patients with Uveal Melanoma Liver Metastases Treated with Isolated Hepatic Perfusion. Ann Surg Oncol 23, 1327–1334 (2016).

17. Chandran, S. S. et al. Treatment of metastatic uveal melanoma with adoptive transfer of tumour-infiltrating lymphocytes: a single-centre, two-stage, single-arm, phase 2 study. Lancet Oncol 18, 792–802 (2017).

18. Forsberg, E. M. V et al. HER2 CAR-T Cells Eradicate Uveal Melanoma and T-cell Therapy-Resistant Human Melanoma in IL2 Transgenic NOD/SCID IL2 Receptor Knockout Mice. Cancer Res 79, 899–904 (2019).

19. Carita, G., Nemati, F. & Decaudin, D. Uveal Melanoma Patient-Derived Xenografts. Ocul Oncol Pathol 1, 161–169 (2015).

20. van der Kooij, M. K., Speetjens, F. M., van der Burg, S. H. & Kapiteijn, E. Uveal Versus Cutaneous Melanoma; Same Origin, Very Distinct Tumor Types. Cancers (Basel) 11, (2019).

21. Durante, M. A. et al. Single-cell analysis reveals new evolutionary complexity in uveal melanoma. Nat Commun 11, 496 (2020).

22. Karlsson, J. et al. Molecular profiling of driver events in metastatic uveal melanoma. Nat Commun 11, 1894 (2020).

23. Rothermel, L. D. et al. Identification of an Immunogenic Subset of Metastatic Uveal Melanoma. Clin Cancer Res 22, 2237–49 (2016).

24. Olofsson, R. et al. Isolated hepatic perfusion as a treatment for uveal melanoma liver metastases (the SCANDIUM trial): study protocol for a randomized controlled trial. Trials 15, 317 (2014).

25. Lin, W., et al. Intra- and intertumoral heterogeneity of liver metastases in a patient with uveal melanoma revealed by single-cell RNA sequencing. Cold Spring Harb Mol Case Stud 7, (2021).

26. Pandiani, C. et al. Single-cell RNA sequencing reveals intratumoral heterogeneity in primary uveal melanomas and identifies HES6 as a driver of the metastatic disease. Cell Death Differ 28, 1990–2000 (2021).

27. Ganesan, A. P. et al. Tissue-resident memory features are linked to the magnitude of cytotoxic T cell responses in human lung cancer. Nat Immunol 18, 940–950 (2017).

28. Li, H. et al. Dysfunctional CD8 T Cells Form a Proliferative, Dynamically Regulated Compartment within Human Melanoma. Cell 176, 775–789.e18 (2019).

29. Huuhtanen, J. et al. Single-cell characterization of anti-LAG3+anti-PD1 treatment in melanoma patients. J Clin Invest (2023) doi:10.1172/JCI164809.

30. Patton, E. E. et al. Melanoma models for the next generation of therapies. Cancer Cell (2021) doi:10.1016/j.ccell.2021.01.011.

31. Einarsdottir, B. O. et al. Melanoma patient-derived xenografts accurately model the disease and develop fast enough to guide treatment decisions. Oncotarget 5, 9609–18 (2014).

32. Jespersen, H. et al. Clinical responses to adoptive T-cell transfer can be modeled in an autologous immune-humanized mouse model. Nat Commun 8, 707 (2017).

33. Ny, L. et al. Supporting clinical decision making in advanced melanoma by preclinical testing in personalized immune-humanized xenograft mouse models. Annals of Oncology **xxx**, (2020).

34. Huuhtanen, J. et al. Evolution and modulation of antigen-specific T cell responses in melanoma patients. Nat Commun 13, (2022).

35. Nilsson, L. M. et al. Genetics and Therapeutic Responses to Tumor-Infiltrating Lymphocyte Therapy of Pancreatic Cancer Patient-Derived Xenograft Models. Gastro Hep Advances 1, 1037–1048 (2022).

36. Ny, L. et al. Supporting clinical decision making in advanced melanoma by preclinical testing in personalized immune-humanized xenograft mouse models. Ann Oncol 31, 266–273 (2020).

37. Ma, D. et al. Patient-derived xenograft culture-transplant system for investigation of human breast cancer metastasis. Commun Biol 4, 1268 (2021).

38. Weiswald, L.-B., Bellet, D. & Dangles-Marie, V. Spherical cancer models in tumor biology. Neoplasia 17, 1–15 (2015).

39. Gopal, S. et al. 3D tumor spheroid microarray for high-throughput, high-content natural killer cell-mediated cytotoxicity. Commun Biol 4, 893 (2021).

40. Al-Hity, G. et al. An integrated framework for quantifying immune-tumour interactions in a 3D co-culture model. Commun Biol 4, 781 (2021).

41. Karlsson, J. et al. Molecular profiling of driver events in metastatic uveal melanoma. Nat Commun 11, 1894 (2020).

42. Chronister, W. D. et al. TCRMatch: Predicting T-Cell Receptor Specificity Based on Sequence Similarity to Previously Characterized Receptors. Front Immunol 12, (2021).

43. Huang, H., Wang, C., Rubelt, F., Scriba, T. J. & Davis, M. M. Analyzing the Mycobacterium tuberculosis immune response by T-cell receptor clustering with GLIPH2 and genome-wide antigen screening. Nat Biotechnol 38, 1194–1202 (2020).

44. Sugase, T. et al. Development and optimization of orthotopic liver metastasis xenograft mouse models in uveal melanoma. J Transl Med 18, 208 (2020).

45. Clarke, J. et al. Single-cell transcriptomic analysis of tissue-resident memory T cells in human lung cancer. Journal of Experimental Medicine 216, 2128–2149 (2019).

46. Fu, Y., Xiao, W. & Mao, Y. Recent Advances and Challenges in Uveal Melanoma Immunotherapy. Cancers (Basel) 14, (2022).

47. van den Berg, J. H., et al. Tumor infiltrating lymphocytes (TIL) therapy in metastatic melanoma: boosting of neoantigen-specific T cell reactivity and long-term follow-up. J Immunother Cancer 8, (2020).

48. Seiter, S. et al. Frequency of MART-1/MelanA and gp100/PMel17-specific T cells in tumor metastases and cultured tumor-infiltrating lymphocytes. Journal of immunotherapy 25, 252–63 (1997).

49. Ye, Q. et al. CD137 accurately identifies and enriches for naturally occurring tumor-reactive T cells in tumor. Clin Cancer Res 20, 44–55 (2014).

50. Inozume, T. et al. Selection of CD8+PD-1+ lymphocytes in fresh human melanomas enriches for tumor-reactive T cells. J Immunother 33, 956–64 (2010).

51. Gros, A. et al. PD-1 identifies the patient-specific CD8^+^ tumor-reactive repertoire infiltrating human tumors. J Clin Invest 124, 2246–59 (2014).

52. Zheng, C. et al. Transcriptomic profiles of neoantigen-reactive T cells in human gastrointestinal cancers. Cancer Cell 40, 410–423.e7 (2022).

53. Simoni, Y. et al. Bystander CD8+T cells are abundant and phenotypically distinct in human tumour infiltrates. Nature 557, 575–579 (2018).

54. Webb, J. R. et al. Profound elevation of CD8+ T cells expressing the intraepithelial lymphocyte marker CD103 (alphaE/beta7 Integrin) in high-grade serous ovarian cancer. Gynecol Oncol 118, 228–36 (2010).

55. Duhen, T. et al. Co-expression of CD39 and CD103 identifies tumor-reactive CD8 T cells in human solid tumors. Nat Commun 9, 2724 (2018).

56. Yee, C. et al. Adoptive T cell therapy using antigen-specific CD8+ T cell clones for the treatment of patients with metastatic melanoma: in vivo persistence, migration, and antitumor effect of transferred T cells. Proc Natl Acad Sci U S A 99, 16168–73 (2002).

57. Dudley, M. E. et al. Cancer regression and autoimmunity in patients after clonal repopulation with antitumor lymphocytes. Science 298, 850–4 (2002).

58. Clémenceau, B. et al. Effector memory alphabeta T lymphocytes can express FcgammaRIIIa and mediate antibody-dependent cellular cytotoxicity. J Immunol 180, 5327–34 (2008).

59. Schreeder, D. M., Pan, J., Li, F. J., Vivier, E. & Davis, R. S. FCRL6 distinguishes mature cytotoxic lymphocytes and is upregulated in patients with B-cell chronic lymphocytic leukemia. Eur J Immunol 38, 3159–66 (2008).

60. Gerlach, C. et al. The Chemokine Receptor CX3CR1 Defines Three Antigen-Experienced CD8 T Cell Subsets with Distinct Roles in Immune Surveillance and Homeostasis. Immunity 45, 1270–1284 (2016).

61. Algazi, A. P. et al. Clinical outcomes in metastatic uveal melanoma treated with PD-1 and PD-L1 antibodies. Cancer 122, 3344–3353 (2016).

62. Rosato, P. C. et al. Virus-specific memory T cells populate tumors and can be repurposed for tumor immunotherapy. Nat Commun 10, 567 (2019).

63. Çuburu, N. et al. Harnessing anti-cytomegalovirus immunity for local immunotherapy against solid tumors. Proc Natl Acad Sci U S A 119, e2116738119 (2022).

64. Hu, Z. I. et al. Immune checkpoint inhibitors unleash pathogenic immune responses against the microbiota. Proc Natl Acad Sci U S A 119, e2200348119 (2022).

65. Fleming, S. J., et al. Unsupervised removal of systematic background noise from droplet-based single-cell experiments using CellBender. *bioRxiv* 791699 (2022) doi:10.1101/791699.

66. Cheloni, S., Hillje, R., Luzi, L., Pelicci, P. G. & Gatti, E. XenoCell: classification of cellular barcodes in single cell experiments from xenograft samples. BMC Med Genomics 14, (2021).

67. Conway, T. et al. Xenome-a tool for classifying reads from xenograft samples. Bioinformatics 28, 172– 178 (2012).

68. Satija, R., Farrell, J. A., Gennert, D., Schier, A. F. & Regev, A. Spatial reconstruction of single-cell gene expression data. Nat Biotechnol 33, 495–502 (2015).

69. Haghverdi, L., Lun, A. T. L., Morgan, M. D. & Marioni, J. C. Batch effects in single-cell RNA-sequencing data are corrected by matching mutual nearest neighbors. Nature Biotechnology 2018 36:5 36, 421–427 (2018).

70. Hippen, A. A. et al. miQC: An adaptive probabilistic framework for quality control of single-cell RNA-sequencing data. PLoS Comput Biol 17, e1009290 (2021).

71. Durante, M. A. et al. Single-cell analysis reveals new evolutionary complexity in uveal melanoma. Nat Commun 11, 496 (2020).

72. Malone, A. F. Monocytes and Macrophages in Kidney Transplantation and Insights from Single Cell RNA-Seq Studies. Kidne*y360* 2, 1654 (2021).

73. Collin, M. & Bigley, V. Human dendritic cell subsets: an update. Immunology vol. 154 3–20 Preprint at 10.1111/imm.12888 (2018).

74. Buonomo, E. L. et al. Liver stromal cells restrict macrophage maturation and stromal IL-6 limits the differentiation of cirrhosis-linked macrophages. J Hepatol 76, 1127–1137 (2022).

75. Zhang, J., He, T., Xue, L. & Guo, H. Senescent T cells: a potential biomarker and target for cancer therapy. EBioMedicine vol. 68 Preprint at 10.1016/j.ebiom.2021.103409 (2021).

76. Zhao, Y., Shao, Q. & Peng, G. Exhaustion and senescence: two crucial dysfunctional states of T cells in the tumor microenvironment. Cellular & Molecular Immunology 2019 17:1 17, 27–35 (2019).

77. Crespo, J., Sun, H., Welling, T. H., Tian, Z. & Zou, W. T cell anergy, exhaustion, senescence, and stemness in the tumor microenvironment. Curr Opin Immunol 25, 214–221 (2013).

78. Ny, L. et al. The PEMDAC phase 2 study of pembrolizumab and entinostat in patients with metastatic uveal melanoma. Nat Commun 12, (2021).

79. Ewels, P. et al. The nf-core framework for community-curated bioinformatics pipelines. (2022) doi:10.5281/ZENODO.7220729.

80. Tickle, T., Tirosh, I., Georgescu, C., Brown, M. & Haas, B. inferCNV of the Trinity CTAT Project. Preprint at https://github.com/broadinstitute/inferCNV (2019).

81. Love, M. I., Huber, W. & Anders, S. Moderated estimation of fold change and dispersion for RNA-seq data with DESeq2. Genome Biol 15, 1–34 (2014).

82. Vita, R. et al. The Immune Epitope Database (IEDB): 2018 update. Nucleic Acids Res 47, D339–D343 (2019).

83. Shugay, M. et al. VDJdb: a curated database of T-cell receptor sequences with known antigen specificity. Nucleic Acids Res 46, D419–D427 (2018).

84. Tickotsky, N., Sagiv, T., Prilusky, J., Shifrut, E. & Friedman, N. McPAS-TCR: A manually curated catalogue of pathology-associated T cell receptor sequences. Bioinformatics 33, 2924–2929 (2017).

85. Gowthaman, R. & Pierce, B. G. TCR3d: The T cell receptor structural repertoire database. Bioinformatics 35, 5323–5325 (2019).

86. Squair, J. W. et al. Confronting false discoveries in single-cell differential expression. Nat Commun 12, (2021).

